# Oncogenic *PIK3CA* promotes cellular stemness in an allele dose-dependent manner

**DOI:** 10.1101/495093

**Authors:** Ralitsa R. Madsen, Rachel G. Knox, Wayne Pearce, Saioa Lopez, Betania Mahler-Araujo, Nicholas McGranahan, Bart Vanhaesebroeck, Robert K. Semple

## Abstract

The *PIK3CA* gene, which encodes the p110α catalytic subunit of PI3-kinase (PI3K), is mutationally activated in cancer and in overgrowth disorders known as *PIK3CA*-related overgrowth spectrum (PROS). To determine the consequences of genetic *PIK3CA* activation in a developmental context of relevance to both PROS and cancer, we engineered isogenic human induced pluripotent stem cells (iPSCs) with heterozygous or homozygous knock-in of *PIK3CA*^*H1047R*^. While heterozygous iPSCs remained largely similar to wild-type cells, homozygosity for *PIK3CA*^*H1047R*^ caused widespread, cancer-like transcriptional remodeling, partial loss of epithelial morphology, upregulation of stemness markers and impaired differentiation to all three germ layers *in vitro* and *in vivo*. Genetic analysis of *PIK3CA*-associated cancers revealed that 64 % had multiple oncogenic *PIK3CA* copies (39 %) or additional PI3K signaling pathway-activating “hits” (25 %). This contrasts with the prevailing view that *PIK3CA* mutations occur heterozygously in cancer. Our findings suggest that a PI3K activity threshold determines pathological consequences of oncogenic *PIK3CA* activation and provide the first insight into the specific role of this pathway in human pluripotent stem cells.

## Significance Statement

The *PIK3CA* H1047R mutation is a common cancer “driver”, and also causes an array of benign but highly disfiguring overgrowth disorders. Human induced pluripotent stem cells engineered to express two copies of *PIK3CA*-H1047R undergo cancer-like transcriptional remodeling and lose their ability to exit the stem cell state. A single mutant copy of *PIK3CA*-H1047R, as observed in non-cancerous overgrowth, had minimal effect on the stem cells and was fully compatible with normal differentiation. Combined with the finding of multiple *PIK3CA* mutant copies in human cancers, this suggests that a signaling threshold determines the disease consequences of *PIK3CA*-H1047R, one of the commonest human oncogenic mutations.

## Introduction

Class IA phosphoinositide 3-kinases (PI3Ks) are essential components of the intracellular signaling cascades triggered by multiple growth factors, especially those acting *via* cell membrane receptor tyrosine kinases. Prominent among these are the insulin and insulin-like growth factor receptors. PI3K signaling is coupled to downstream activation of AKT and mammalian target of rapamycin complex 1 (mTORC1), which play key roles in organismal growth and development (1–3).

Strongly kinase-activating mutations in *PIK3CA*, the gene encoding the catalytic p110α subunit of PI3K, are among the most frequently observed oncogenic events in a range of human tumors (4). Although widely referred to as cancer “drivers”, the same mutations have also been identified in non-malignant, albeit often severe, overgrowth disorders (5). These disorders are caused by postzygotic mosaic *PIK3CA* mutations and are phenotypically diverse, reflecting different patterns of mutation distribution and likely also different strengths of PI3K activation.

The commonest *PIK3CA* “hotspot” variant, H1047R, has been studied extensively in cancer models, both in cells and *in vivo*. Endogenous, heterozygous expression in mice seemingly only results in cancer development in combination with additional oncogenic drivers (6–11), while transgenic overexpression of this *PIK3CA* mutant does lead to early malignancy (12–17). In cultured cells, *PIK3CA*^H1047R^ overexpression, but not heterozygous expression from the endogenous locus, leads to cellular transformation (18, 19). In human tumors, *PIK3CA* mutations are not mutually exclusive with other oncogenic alterations within the PI3K pathway (20), suggesting that stronger pathway activation may be required for malignant progression. This is supported by the benign nature of the overgrowth in PROS where *PIK3CA*^H1047R^ heterozygosity is not sufficient to cause cancer. Despite this circumstantial evidence of dose-dependent effects of genetic PI3K activation, this has not been examined directly owing to the paucity of isogenic experimental models with endogenous expression of a defined number of oncogenic variants.

Disorders such as PROS illustrate that understanding aberrant development may hold lessons for cancer (21). Malignant transformation of cells typically involves dedifferentiation, reactivation of developmental pathways and phenotypic plasticity. *PIK3CA*^H1047R^ was recently linked to induction of multipotency and cellular dedifferentiation in two mouse models of breast cancer (8, 16). Overexpression of wild-type (WT) *PIK3CA* in the head and neck epithelium of a mouse model of oral carcinogenesis has also been associated with dedifferentiation and epithelial-to-mesenchymal transition, increased transforming growth factor β (TGFβ) signaling and upregulated expression of the pluripotency factors *Nanog* and *Pou5f1* (*Oct3/4*) (22). Despite the insights gained from these and other mouse models of oncogenic *PIK3CA*, efforts to establish *in vivo* models of PROS have highlighted that species differences may constrain extrapolation from model organisms to the mechanisms of pathological PI3K activation in human disease (5).

Due to their unlimited self-renewal and differentiation capacity, human pluripotent stem cells are increasingly used as tools to develop more relevant human disease models (23). Their inherent similarities to cancer cells also make them an attractive system in which to study oncogenic processes (24). Thus, to study dose-dependent effects of pathological PI3K hyperactivation in a developmental system of relevance to cancer and PROS, we engineered isogenic human induced pluripotent stem cells (iPSCs) to express *PIK3CA*^H1047R^ from one or both endogenous loci. Our data reveal clear dose-dependent developmental phenotypes downstream of p110α activation, with homozygosity but not heterozygosity for *PIK3CA*^H1047R^ promoting self-sustained stemness *in vitro* and *in vivo*. These findings emphasize the importance of using precisely-engineered models of cancer-associated *PIK3CA* variants to obtain a faithful representation of their biological effects and have implications for our understanding of PI3K activation in human disease.

## Results

### Generation of human iPSCs with endogenous expression of *PIK3CA H1047R*

To establish a cell model suitable for interrogation of allele dose-dependent consequences of p110α activation in human development and disease, we used CRISPR/Cas9 genome editing of well-characterized, karyotypically normal wild-type (WT) iPSCs to generate multiple isogenic clones either heterozygous (*n* = *3*) or homozygous (*n* = *10*) for the activating *PIK3CA*^H1047R^ allele (**Figure 1a,b**, **Figure S1a**). To control for non-specific effects caused by genetic drift following so-called bottleneck selection (25, 26), we expanded six WT clones exposed to the gene targeting process. Sequencing of multiple clones of each genotype showed no evidence of mutagenesis of 17 computationally predicted CRISPR off-target sites (**Figure S1b**), and a normal karyotype was confirmed in three homozygous and two heterozygous clones more than 10 passages after targeting (**Figure S1c**), suggesting that *PIK3CA*^H1047R^ does not lead to widespread genomic instability in these cells.

**Figure 1.**
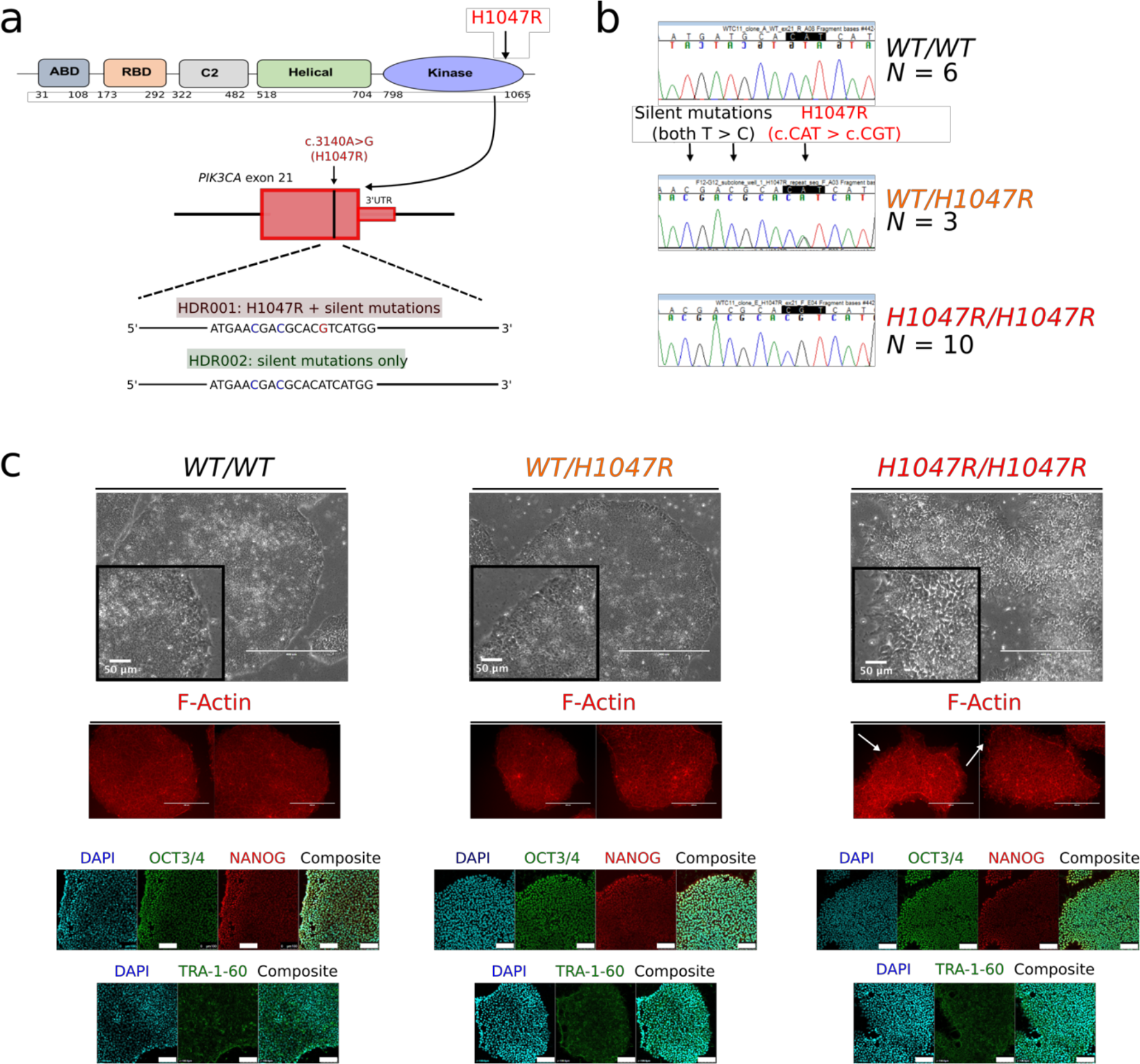
Generation of isogenic human pluripotent stem cells expressing *PIK3C4*^H1047R^ from one or both endogenous alleles. **a.** Domain structure of the *PIK3CA* gene product p110α and schematic of the CRISPR/Cas9 targeting strategy showing the two homology-directed repair (HDR) templates used alone or in combination. **b.** Representative sequences of heterozygous and homozygous *PIK3CA*^*H1047R*^ clones, all homozygous for the two silent mutations introduced by the targeting strategy. The number of independent clones of each genotype generated is provided next to the chromatograms. WT: wild-type. **c.** Representative light microscopy and immunofluorescence images of stem cell colonies from cultures with the indicated genotypes. F-Actin staining was used to visualize cell shape, and arrows highlight altered edge morphology and F-Actin-rich protrusions in *PIK3CA*^*H1047R/H1047R*^ colonies. Scale bars = 400 μm (50 μm for inserts). Images are representative of all clones of the corresponding genotype. The confocal images are of wild-type and mutant cells stained with antibodies against OCT3/4, NANOG and TRA-1-60. Images are representative of at least 2 independent experiments and clones per genotype. Scale bar = 100 μm. See also Figure S1.

WT and *PIK3CA*^*WT/H1047R*^ colonies had a similar microscopic appearance, whereas *PIK3CA*^*H1047R/H1047R*^ clones exhibited aberrant colony morphology, characterized by disorganization of the normal epithelial appearance, including pronounced F-Actin-rich protrusions visible on colony margins (**Figure 1c**). Homozygous cells also proved more adherent in routine passaging, requiring longer dissociation time than WT and heterozygous cultures. Nevertheless, *PIK3CA*^*H1047R/H1047R*^ clones remained positive for the pluripotent stem cell markers NANOG, OCT3/4 and TRA-1-60 (**Figure 1c**), consistent with preserved stem cell identity.

### Allele dose-dependent signaling effects of *PIK3CA*^*H1047R*^

We next assessed PI3K signaling in *PIK3CA*^*wt/H1047R*^ and *PIK3CA*^*H1047R/H1047R*^ iPSCs. p110α protein expression was reduced in both mutant genotypes and sometimes barely detectable in *pIK3CA*^*H1047R/H1047R*^ cells. Despite this, immunoblotting revealed graded increases in AKT phosphorylation across *PIK3CA*^*WT/H1047R*^ and *PIK3CA*^*H1047R/H1047R*^ lines, both in growth factor-replete conditions (**Figure 2a**) and upon growth factor removal (**Figure 2b**). Consistent with previous findings in breast epithelial cells heterozygous for *PIK3CA*^H1047R^ (19), both *PIK3CA*^*WT/H1047R*^ and *PIK3CA*^*H1047R/H1047R*^ cells also showed modest and graded increases in ERK phosphorylation.

**Figure 2.**
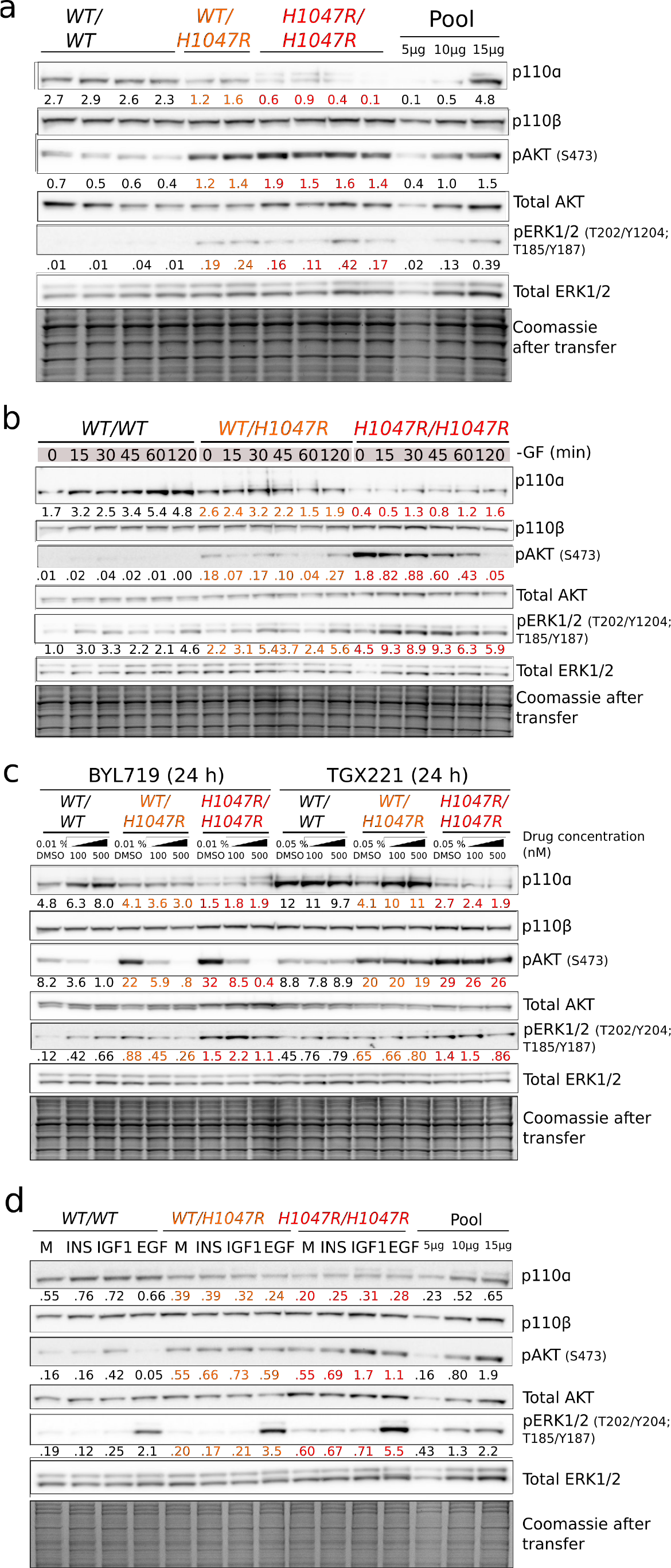
Graded activation of PI3K signaling in *P1K3CA*^H1047R^ human pluripotent stem cells. Immunoblots are shown for p110α and p110β catalytic subunits of PI3K, and for total and phosphorylated AKT and ERK, with Coomassie-stained gels after transfer as a control. Numbers below bands indicate quantification by densitometry (arbitrary units). **a.** Signaling in cells collected 3 h after replenishment of growth factor (GF)-replete medium. Representative of ≥3 independent experiments. **b.** Signaling time course during short-term GF depletion. Representative of≥ 2 independent experiments. **c.** Effects of 24 h of specific p110α or p110β inhibition in GF-replete medium using BYL719 or TGX221, respectively. DMSO was used as control. Representative of 2 independent experiments. See also Figure S2. **d.** Response of cells to 2 h of GF depletion followed by 20 min stimulation with 10 nM of insulin (INS), insulin-like growth factor 1 (IGF1) or epidermal growth factor (EGF). GF-free DMEM/F12 medium (M) was used as control. The results are representative of 2 independent experiments. In all cases independent clones of the same genotypes were used for replicate experiments. Protein pool dilutions are included where possible to assess assay performance.

Baseline PI3K pathway hyperactivation was inhibited in a dose-dependent manner by the p110α-specific inhibitor BYL719, while the p110β-specific inhibitor TGX221 had no effect (**Figure 2c**). BYL719 did not reverse the allele dose-dependent downregulation of the p110α protein, suggesting that it is not caused by acute negative feedback mechanisms. In both mutant genotypes, low-dose BYL719 (100 nM) reduced AKT phosphorylation to the level in untreated WT cells (**Figure 2c**), without inhibiting growth (**Figure S2a**). Relative to WT controls, mutant stem cells exhibited increased survival upon prolonged growth factor depletion, and this was also reversed by low-dose BYL719 (**Figure S2b**). A higher concentration of BYL719 (500 nM) was cytotoxic to both WT and *PIK3CA*^*WT/H1047R*^ cells (**Figure S2a**), but not *PIK3CA^H1047R/H1047R^* cells, in which it reversed the aberrant colony morphology (**Figure S2a,c**).

We also examined responses to acute stimulation with insulin, insulin-like growth factor 1 (IGF1) or epidermal growth factor (EGF) (**Figure 2d**). *PIK3CA*^*WT/H1047R*^ and *PIK3CA*^*H1047R/H1047R*^ cells had high baseline AKT phosphorylation. This exceeded the level in IGF1-stimulated WT cells, but no consistent increase in the response to IGF 1 was seen in mutant cells compared to WT (**Figure 2d**). Insulin did not elicit discernible AKT phosphorylation in any of the iPSC cells used. This apparent insulin resistance may be caused by the high concentration of insulin (3 μM) used in the maintenance medium (27), resulting in downregulation of insulin receptor expression at the plasma membrane (28). A modest increase in AKT phosphorylation in response to EGF was only observed in homozygous mutant cells. In contrast, EGF stimulation enhanced ERK phosphorylation above baseline in all iPSC lines, and this was progressively enhanced across heterozygous and homozygous mutant cells (**Figure 2d**). These findings suggest that the MAPK/ERK pathway is primed to hyper-respond to growth factor stimulation in *PIK3CA*^*H1047R*^ stem cells, in an allele dose-dependent manner.

### Transcriptomic effects of *PIK3CA*^H1047R^ in pluripotent stem cells

To determine the wider dose-dependent consequences of genetic p110α activation, we profiled the protein-coding transcriptome of WT, *PIK3CA*^*WT/H1047R*^ and *PIK3CA^H1047R/H1047R^* iPSCs, cultured in growth factor-replete conditions to mimic the *in vivo* milieu of the pluripotent epiblast. Multidimensional scaling demonstrated distinct transcriptomic signatures of WT, heterozygous and homozygous cells (**Figure 3a**). The transcriptome of *PIK3CA*^*WT/H1047R*^ cells was nearly identical to WT controls, with only 131 differentially-expressed transcripts (FDR = 0.05). In contrast, homozygosity for *PIK3CA*^H1047R^ led to differential expression of 1,914 genes (**Figure 3a**). This indicates widespread transcriptional remodeling with a sharp allele dose-dependency, suggestive of a threshold effect.

**Figure 3.**
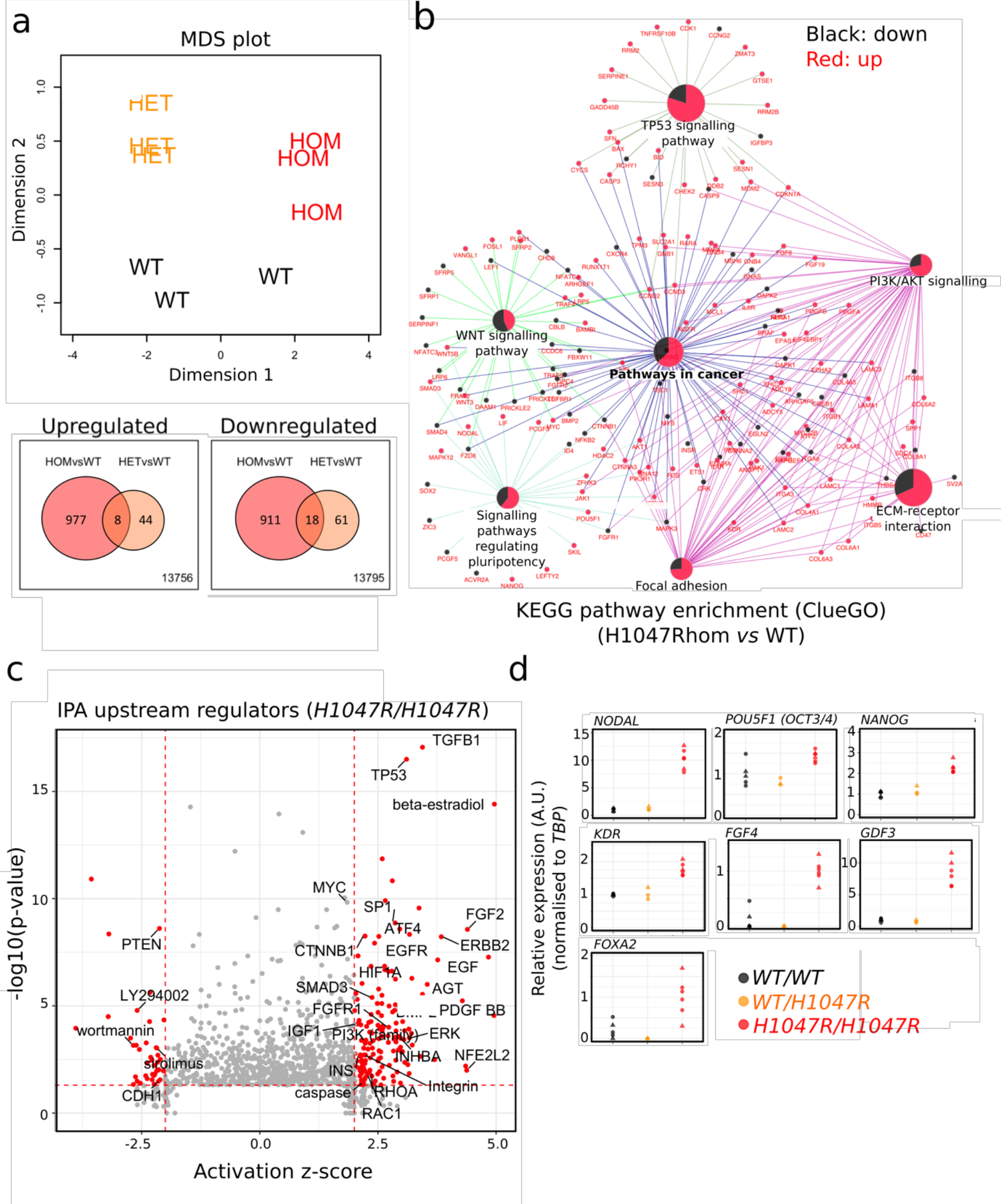
Widespread transcriptional remodeling in *PIK3CA*^*H1047R/H1047R*^ pluripotent stem cells. **a.** Top: Multidimensional scaling (MDS) plot of transcriptomes of wild-type (WT), *PIK3CA*^*WT/H1047R*^ and *PIK3CA*^*H107R/H1047R*^ human pluripotent stem cells profiled by RNAseq. Bottom: Venn diagrams showing overlap of upregulated and downregulated transcripts in *PIK3CA*^*H1047R*^ mutants compared to WT (FDR < 0.05; Benjamini-Hochberg; 3 clones per genotype). FC, fold-change. **b.** KEGG Pathway Enrichment Analysis undertaken using ClueGO on the 1,914 differentially regulated transcripts in *PIK3CA*^*H1047R/H1047R*^ iPSCs. Pathways shown were significantly enriched with FDR=0.05 (Benjamini-Hochberg). Differentially expressed genes belonging to enriched pathways are shown in red (upregulated) or black (downregulated). The proportions of upregulated and downregulated genes within a pathway are represented in central pie charts. ECM, extracellular matrix. **c.** Ingenuity^®^ Pathway Analysis (IPA) of upstream regulators in *PIK3CA*^*H1047R/H1047R*^ cells, based on all differentially expressed genes. Components with absolute activation z-score > 2 and p-value < 0.05 are highlighted in red. Selected components linked to PI3K signaling and pluripotency are labelled. **d.** Assessment of expression of selected epiblast genes based on the pathway analyses in 4b and 4c. Data were obtained in 2 independent experiments. Expression values were scaled to the WT or *PIK3CA*^*H1047R/H1047R*^ mean as indicated. Individual points correspond to separate cultures: 5 WT (3 clones), 3 *PIK3CA*^*WT/H1047R*^ (2 clones) and 6 *PIK3CA*^*H1047R/H1047R*^ (4 clones). All clones used for confirmation were distinct from those used to generate RNAseq data. See also Figure S3.

KEGG annotation-based pathway analysis using all 1,914 differentially-expressed genes in *PIK3CA*^*H1047R/H104m7R*^ cells demonstrated significant changes to PI3K/AKT signaling, as expected. “Pathways in cancer” was identified as a common central node, demonstrating the power of our isolated genetic activation of PI3K to recapitulate signatures identified in the genetically far more chaotic context of tumors (**Figure 3b**). Other pathways identified as showing coherent perturbations were “Extracellular matrix-receptor interaction” and “Focal adhesion”, in keeping with the altered morphology and adhesion properties of homozygous mutants. Several genes involved in pluripotency regulation and WNT signaling were also differentially expressed. Finally, the TP53 pathway was found to be significantly altered (**Figure 3b**). This is consistent with prior evidence of TP53 activation in cell lines with hyperactivation of PI3K/AKT (29–32), however given the recent report that a substantial proportion of iPSC lines have *TP53* mutations (33), we sequenced the *TP53* gene of all clones. We found that two of the WT lines were indeed heterozygous for *TP53* C135F (**Figure S3a**), a mild loss-of-function allele based on biochemical assays in yeast (34). Despite this, inspection of each iPSC clone’s RNAseq data for the differentially expressed TP53 signaling genes showed that the signature difference in *PIK3CA*^*H1047R/H1047R*^ cells was not attributable to these two WT lines.

To identify potential drivers of the transcriptional changes in *PIK3CA*^*H1047R/H1047R*^ cells, we also undertook Ingenuity^®^ Pathway Analysis of upstream regulators. This again revealed the expected activation of PI3K/AKT signaling. It also implicated factors important in stem cell regulation, including TGFβ, FGF2, TP53, β-catenin and MYC (**Figure 3c**). TGFβ was the most significant prediction, and supporting increased signaling within this pathway, we found increased phosphorylation of SMAD2 in homozygous mutants (**Figure S3b**). These cells also had upregulated expression of *NODAL* (**Figure 3b, d**), a member of the TGFβ superfamily that maintains the pluripotent epiblast at early developmental stages and later induces primitive streak formation during gastrulation (35). Consistent with NODAL’s dual function, *PIK3CA*^*H1047R/H1047R*^ cells exhibited a stemness signature (36) including upregulation of *NANOG*, *POU5F1* (*OCT3/4*), *MYC*, *KDR*, *IGF1R*, as well as upregulation of primitive streak markers such as *FGF4*, *GDF3* and *FOXA2* (**Figure 3b, d**). Upregulation of *NODAL* in WT and mutant cells was abolished by p110α-specific inhibition with BYL719 (**Figure S3c**). In comparison, *NANOG* expression remained mostly unaffected by BYL719, with a trend towards downregulation after 48 h of p110α inhibition (**Figure S3c**). These findings suggest upregulation of *NODAL* and enhanced TGFβ/SMAD2 signaling as a candidate mechanism whereby p110α activation may exert effects on stemness in human pluripotent stem cells.

### Homozygosity for *PIK3CA*^H1047R^ confers self-sustained stemness upon embryoid bodies

Embryoid bodies (EBs) are widely used to model lineage specification during gastrulation (37, 38). Previous studies have shown that *NODAL* overexpression in human pluripotent stem cell-derived EBs blocks differentiation to all three germ layers (39). Given the evidence for upregulated *NODAL* and TGFβ signaling in *PIK3CA*^*H1047R/H1047R*^ cells, we tested whether the resulting EBs would behave similarly to NODAL-overexpressing EBs. EBs were established without TGFβ and FGF2, cultured in suspension for four days and allowed to generate adherent outgrowths for six days (**Figure 4a**). *PIK3CA*^*H1047R/H1047R*^ stem cells consistently generated compact, cystic EBs that failed to bud and undergo internal reorganization (**Figure 4b**), with notable resemblance to mouse EBs overexpressing constitutively-active PDK1 or AKT1 (40). In adherent culture, *PIK3CA*^*H1047R/H1047R*^ EB outgrowths resembled stem cell colonies (**Figure 4b**). Confirming this, *PIK3CA*^*H1047R/H1047R*^ EB outgrowths stained positive for the stemness markers OCT3/4, NANOG and TRA-1-60 (**Figure 4c**). WT and *PIK3CA*^*WT/H1047R*^ EBs, in contrast, exhibited complex morphologies in suspension and yielded heterogeneous outgrowths of differentiated cells which continued to mature during the experiment (**Figure 4b,c**).

**Figure 4.**
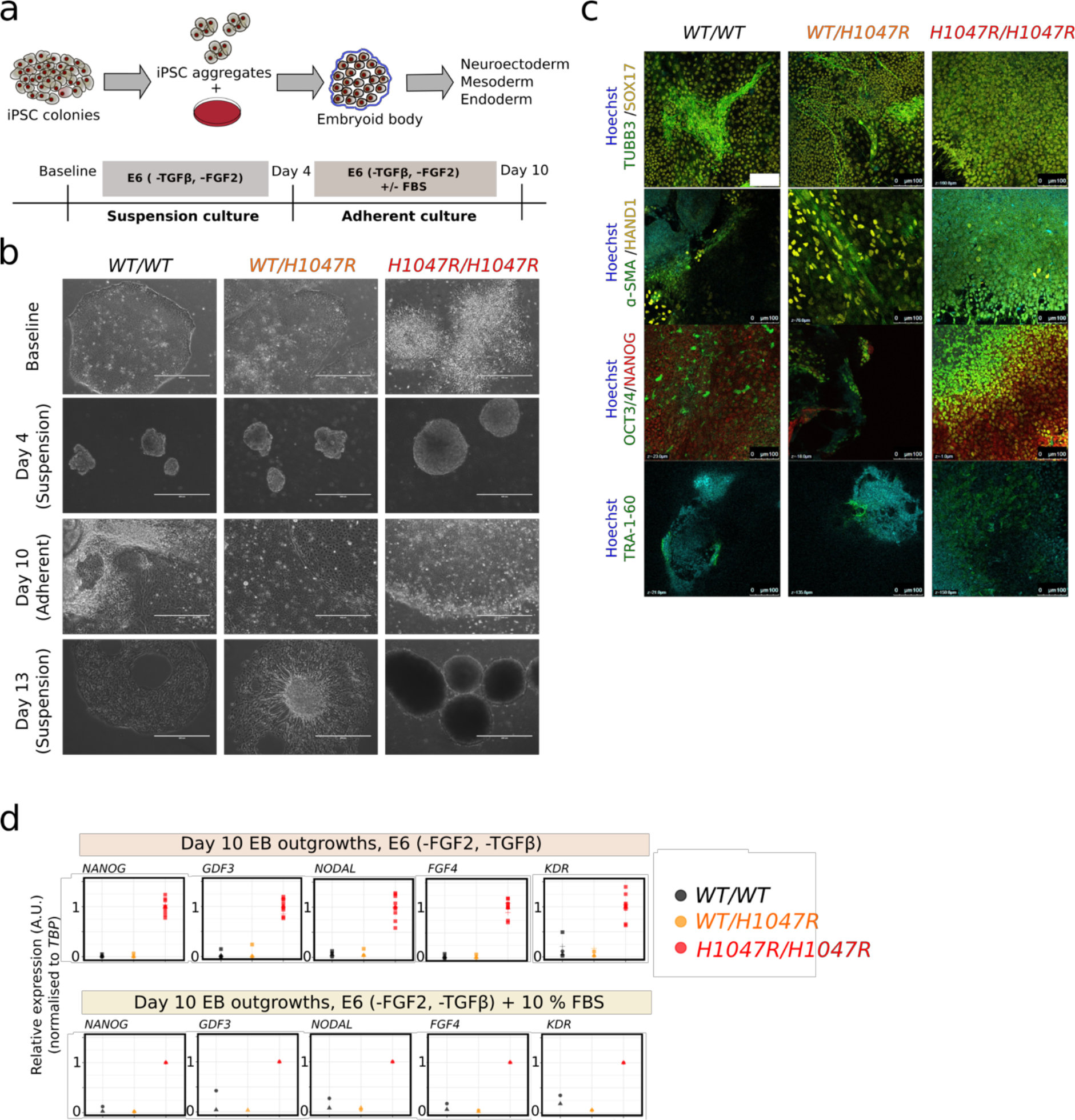
Self-sustained stemness in *PIK3CA*^*H1047R/H1047R*^ embryoid bodies. **a.** Schematic illustrating the protocol used for embryoid body (EB) formation and subsequent adherent culture. E6, Essential 6 medium; FBS, fetal bovine serum; FGF2, fibroblast growth factor 2; TGFβ, transforming growth factor β. **b.** Representative brightfield micrographs of wild-type (*WT/WT*), *PIK3CA*^*WT/H1047R*^ and *PIK3CA*^*H1047R/H1047R*^ cells at baseline (iPSC stage), 4 days (suspension), 10 days (adherent) and 13 days (suspension) following EB formation. *PIK3CA*^*H1047R/H1047R*^ iPSC colonies are refiactile due to partial dissociation, while stem cell-like colonies emerging from adherent *PIK3CA*^*H1047R/H1047R*^ EBs are highly compact. In addition to the floating layers of differentiated cells shown here, WT and *PIK3CA*^*WT/H1047R*^ suspension cultures on day 13 also contained larger EB aggregates with complex morphologies and internal differentiation. Scale bar = 400 μm. **c.** EB outgrowths were fixed on day 10 and stained for TRA-1-60 or co-stained for TUBB3/SOX17, β-SMA/HAND1 or NANOG/OCT3/4. Hoechst was used for nuclear visualization. Images are representative of 2 independent experiments, using a single WT clone and 2 clones of each mutant. Scale bar = 100 μm. **d.** Real time quantitative PCR analysis of stemness gene expression in EB outgrowths in E6 medium without TGFβ and FGF2. Individual replicates shown in the panel are from 3-4 WT clones, 2 *PIK3CA*^*WT/H1047R*^ clones and 4 *PIK3CA*^*H1047R/H1047R*^ clones. Duplicate outgrowth cultures of *PIK3CA*^*H1047R/H1047R*^ clones are also shown. Expression values are in arbitrary units (A.U.). See also Figure S4.

The apparent differentiation block of *PIK3CA*^*H1047R/H1047R*^ EBs was assessed transcriptionally using lineage-specific arrays and candidate gene quantitative PCR. Unlike WT and *PIK3CA*^*WT/H1047R*^ EBs, homozygous mutants exhibited sustained expression of stemness genes and failed to upregulate germ layer-specific markers, both in adherent cultures and in suspension (**Figure 4d**, **Figure S4a-d**). This phenotype persisted in the presence of serum, which is used to induce EB differentiation (**Figure 4d**, **Figure S4a**). Attempts to reverse the *PIK3CA*^*H1047R/H1047R*^ EB phenotype with the p110α inhibitor BYL719 were unsuccessful due to poor EB survival in the presence of the drug, consistent with previous studies demonstrating high EB sensitivity to PI3K/mTOR inhibition (40–42).

### Heterozygosity for *PIK3CA*^H1047R^ is compatible with directed definitive endoderm formation

Heterozygosity for *PIK3CA*^*H1047R*^ did not produce major perturbations in the transcriptome of iPSCs nor in EB differentiation. Nevertheless, observation of *PIK3CA*-driven overgrowth in PROS suggests that mesodermal and neuroectodermal tissues are widely involved while tissues of endodermal origin are only rarely affected by strong activating mutations, raising the possibility of negative selection during endodermal development (5). We thus sought to undertake more systematic analysis of early differentiation in our human developmental models of *PIK3CA*^*H1047R*^. To overcome the high variability seen in self-aggregating, spontaneously-differentiating EBs, the protocol was modified (**Figure 5a**), incorporating use of microwell plates to ensure homogeneous EB size. EB formation was followed by three days of exposure to different concentrations of Activin A, BMP4 and FGF2 to promote mesoderm or definitive endoderm formation (43, 44). Lineage-specific gene expression arrays, candidate gene quantitative PCR and immunostaining assays were used to assess expression of multiple differentiation markers (**Figure 5b,c**). Mesoderm or endoderm induction led to increased expression of the expected lineage-specific markers (**Figure 5b**, **Figure S5**). The temporal pattern and relative expression levels of the analyzed genes was similar for *PIK3CA*^*WT/H1047R*^ and WT EBs (**Figure 5b**, **Figure S5**), and adherent outgrowths from both stained positive for mesoderm and endoderm markers at the end of the 10-day protocol (**Figure 5c**). The results of this assay argue against an inability of *PIK3CA*^*WT/H1047R*^ iPSCs to yield definitive endoderm.

**Figure 5.**
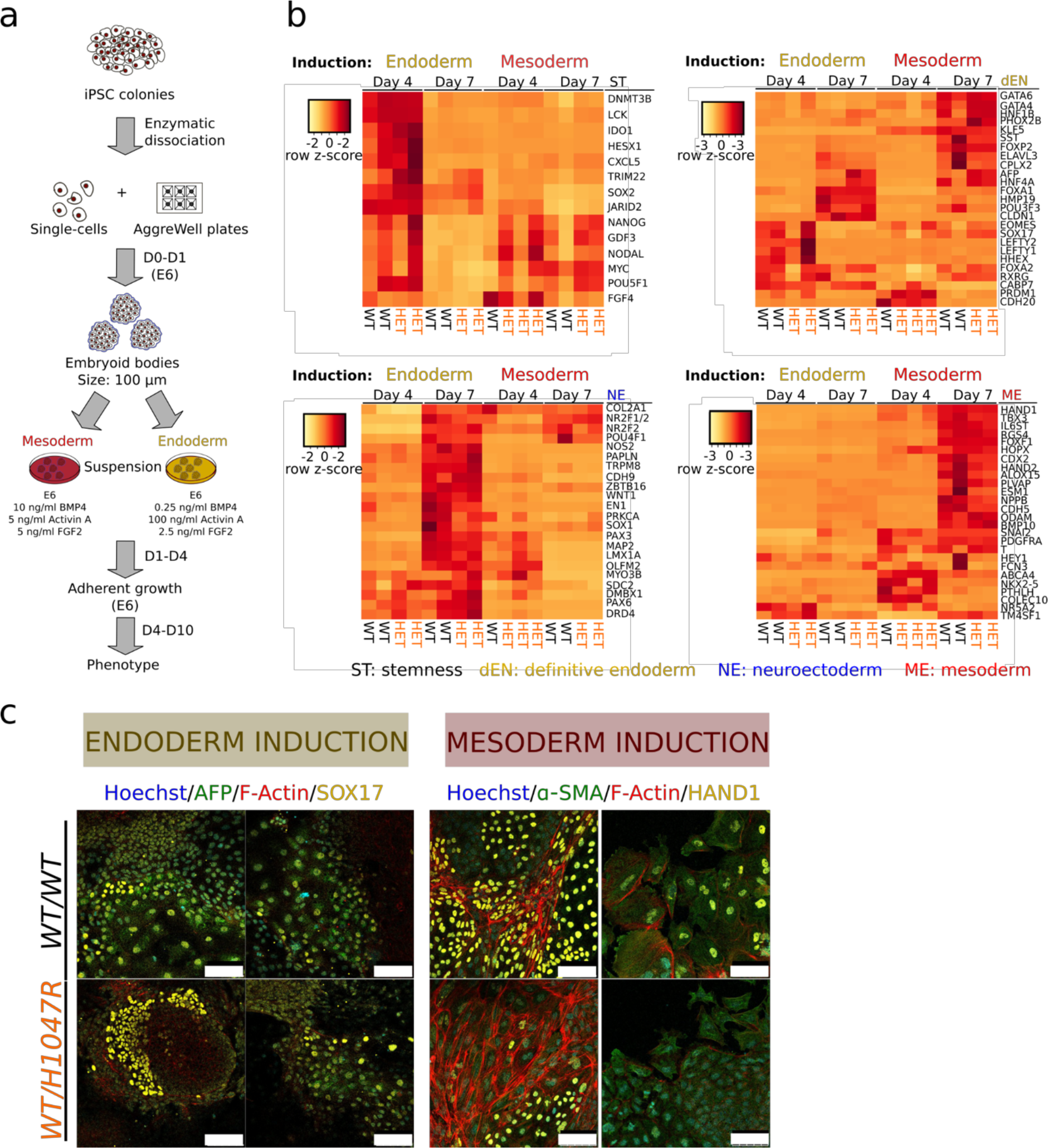
Heterozygosity for *PIK3CA*^H1047R^ does not affect endoderm or mesoderm differentiation in embryoid bodies. **a.** Schematic illustrating the AggreWell-based protocol for embryoid body (EB) culture with subsequent 3-day mesoderm or endoderm differentiation in suspension culture, followed by transfer to adherent conditions. D, day; E6, Essential 6 medium. **b.** Real time quantitative PCR scorecard-based profiling of lineage markers in wild-type (WT) and *PIK3CA*^*WT/H1047R*^ EBs following mesoderm or endoderm induction. Gene heatmaps are shown across rows and are grouped according to lineage. Colors correspond to expression z-scores as indicated. “Endoderm” and “Mesoderm” indicate induction conditions. Results are from a single experiment examining the following replicates: day 4 endoderm: 2 WT clones and 2 cultures of a single H1047Rhet clone; day 4 mesoderm: 1 WT clone and 3 H1047Rhet cultures from 2 clones; day 7 endoderm and mesoderm: 2 WT clones and 2 separate cultures of a single H1047Rhet clone. See also Figure S5 for RT-qPCR validation and additional replicates, including gene expression analysis on day 10. **c.** Representative confocal images of WT and H1047Rhet Eb outgrowths on day 10 of the differentiation protocol, stained with antibodies against endoderm (AFP/SOX17) and mesoderm (α-SMA/HAND1)-specific markers. Hoechst was used for nuclear visualization and F-Actin to for cell boundary demarcation. The images are from one clone per genotype. Scale bar = 100 μm.

We also subjected WT and *PIK3CA*^*H1047R*^-harbouring cell lines to monolayer-based directed differentiation using a combination of low serum, inhibition of GSK3 and high levels of Activin A (45) (**Figure 6a**). The differentiation medium was also supplemented with DMSO (control) or BYL719 (100 nM), in anticipation that high PI3K signaling would be incompatible with two-dimensional definitive endoderm formation, as reported previously (46, 47). Unexpectedly, both *PIK3CA*^*WT/H1047R*^ and *PIK3CA*^*H1047R/H1047R*^ iPSCs differentiated successfully to definitive endoderm under these directed conditions, as evidenced by gene expression analysis and immunostaining (**Figure 6b,c**). The dynamics of gene expression were closely similar across the three genotypes and were unaffected by p110α inhibition (**Figure 6b**). Confirming that this was not a donor-specific effect, similar results were obtained with isogenic WT and mutant iPSCs derived from a PROS patient with mosaic, heterozygous expression of the rare *PIK3CA*^*E418K*^ allele (**Figure S6**). These findings suggest that PI3K activation is compatible with definitive endoderm formation *in vitro*, contrary to previous conclusions based on the use of non-specific pan-PI3K inhibitors with known off-target effects (46, 47), and do not support cell-autonomous negative selection in early lineage specification in PROS.

**Figure 6.**
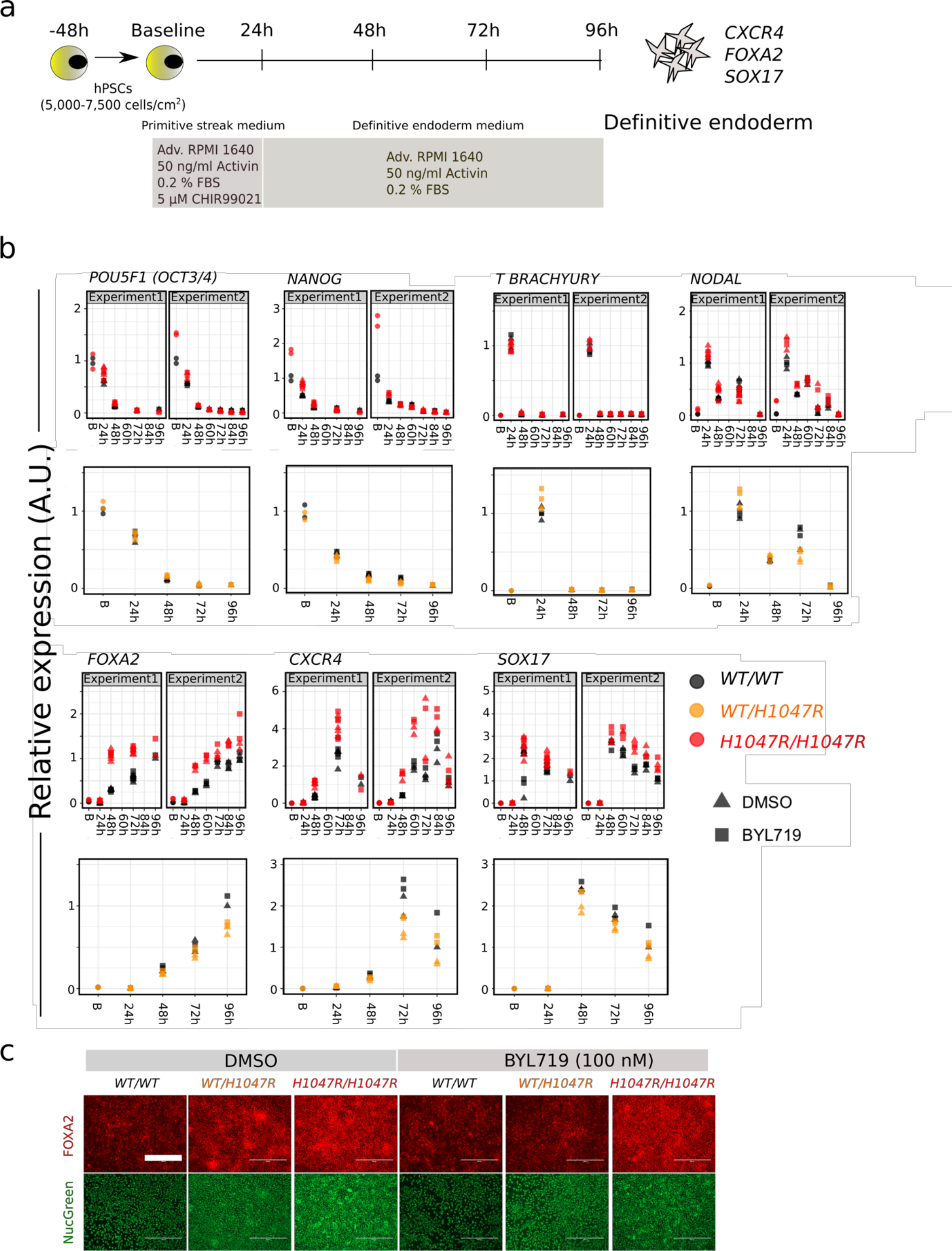
*PIK3C4*^H1047R^ is compatible with monolayer definitive endoderm differentiation. **a.** Schematic of the protocol for definitive endoderm differentiation in monolayer culture. Adv., advanced; FBS, fetal bovine serum. **b.** Real time quantitative PCR analysis of lineage marker expression during differentiation in the presence of DMSO (control) or the p110α-specific inhibitor BYL719 at 100 nM. Data from 2 independent experiments with WT *vs PIK3CA*^*H1047R/H1047R*^ are shown side by side. 2 cultures of each of 2 clones per genotype were profiled. The time course data for *PIK3CA*^*WT/H1047R*^ vs WT cells are from a single experiment using 2 cultures of 1 clone per genotype. Gene expression was scaled internally to the mean value of an appropriate time-point, and resulting values are arbitrary. See also Figure S6. **c.** Immunofluorescence staining for FOXA2 in WT, H1047Rhet and *PIK3CA*^*H1047R/H1047R*^ definitive endoderm cultures at the end of differentiation. NucGreen was used for nuclear visualization. Note the higher cell density in *PIK3CA*^*H1047R/H1047R*^ cultures, attributable to improved survival following single-cell passaging. Scale bar=400 μm.

### Allele dose-dependent effects of *PIK3C4*^H1047R^ *in vivo*

To confirm that allele dose-dependent effects of *PIK3CA*^*H1047R*^ were not artefacts of *in vitro* culture, we injected immunodeficient mice with WT or mutant iPSCs, and allowed them to form tumors over 5-8 weeks before histopathological assessment. WT and *PIK3CA*^*WT/H1047R*^ tumors contained differentiated components of the three germ layers, including bone, cartilage, pigmented epithelium, nervous tissue and tubular endodermal structures (**Figure 7a**, **Table S1**). All *PIK3CA*^*WT/H1047R*^ tumors exhibited better differentiated endoderm-derived tissues including respiratory (all lines) and gastrointestinal (one line) epithelium, corroborating the *in vitro* finding that heterozygosity for *PIK3CA*^*H1047R*^ does not impair definitive endoderm formation. In contrast, differentiated components were either completely absent or very immature in the two *PIK3CA*^*H1047R/H1047R*^ tumors (**Figure 7a**, **Table S1**), consistent with the inability of the parental cells to yield spontaneously-differentiated EBs. The least mature of the *PIK3CA*^*H1047R/H1047R*^ tumors showed extensive recruitment of mouse stromal cells, forming septae separating lobules of immature human tissue (**Figure S7a**). Homozygous tumors also exhibited extensive necrosis and yolk sac-like tissue formation (**Figure 7a**), the latter suggested to be an *in vivo* characteristic of injected pluripotent stem cells with malignant potential (48). They also contained multiple foci positive for T BRACHYURY (immature mesoderm) and nuclear OCT3/4 (embryonal carcinoma marker in germ cell tumors) (**Figure S7c,d**).

These results are in line with the *in vitro* studies and demonstrate that homozygosity but not heterozygosity for *PIK3CA*^*H1047R*^ promotes stemness of human pluripotent stem cells. Stem cells share many similarities with cancer cells, and phenotypes such as dedifferentiation and reactivation of developmental pathways have been linked to epithelial-to-mesenchymal transition and aggressive tumor behavior *in vivo* (49). *PIK3CA* mutations in human tumors are not mutually-exclusive with other oncogenic alterations promoting PI3K pathway activation, suggesting that further activation is positively selected for (50). This raises the possibility that our findings may be relevant to understanding the behavior of human cancer. We thus analyzed the prevalence of multiple oncogenic “hits” within the PI3K pathway using data from The Cancer Genome Atlas on cancer types with >2014;10% prevalence of *PIK3CA* mutations. In aggregate, 21% of these cancers had *PIK3CA* mutations. Nearly 40% of this subset had more than one copy of the mutation, and 25% also had a mutation in other selected PI3K pathway components (*PTEN, PIK3R1, AKT1/2/3*), or harbored a second *PIK3CA* variant (**Figure 7b**). This high frequency of additional mechanisms activating PI3K signaling in cancers provides circumstantial support for the notion that the strength of PI3K hyperactivation may be important for tumor progression *in vivo*.

**Figure 7.**
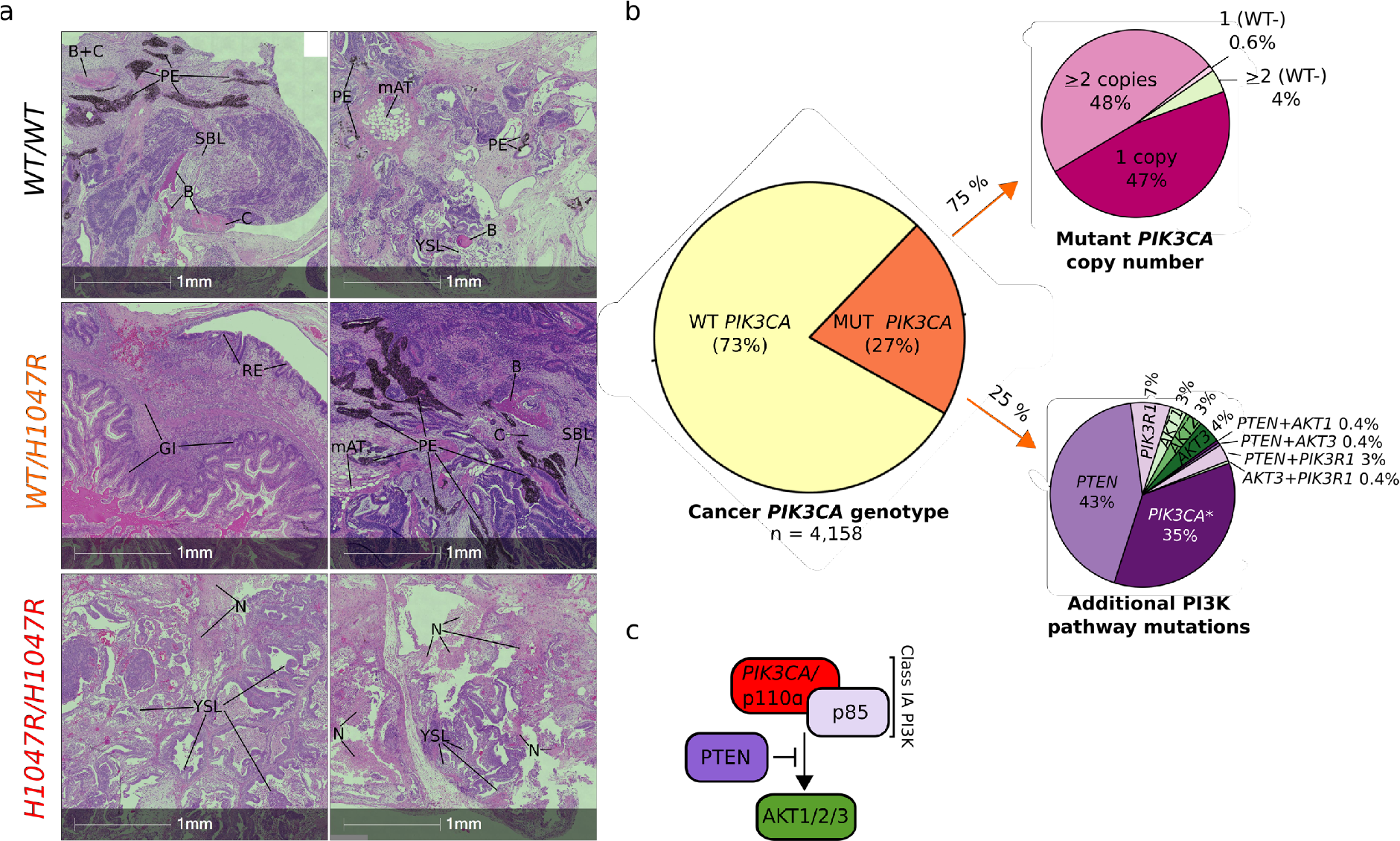
*PIK3C4*^H1047R^ allele dose-dependent effects in tumor xenografts and genetic evidence for graded PI3K activation in cancers. **a.** Hematoxylin and eosin-stained (H&E) sections of WT, *PIK3CA*^*WT/H1047R*^ and *PIK3CA*^*H1047R/H1047R*^ tumor xenografts derived from injection of human pluripotent stem cells into immunodeficient mice. The micrographs are from 2 tumors per genotype and are representative of a total of 5, 3 and 2 tumors from WT, *PIK3CA*^*WT/H1047R*^ and *PIK3CA*^*H1047R/H1047R*^ respectively. Yolk sac-like (YSL) and embryonal carcinoma-like (ECL) tissues, suggesting neoplastic transformation of cells within the original cultures, were more prevalent in *PIK3CA*^*H1047R/H1047R*^ tumors, which also exhibited extensive necrosis (N); rare YSL foci were seen in two other tumors derived from the same WT clone. The only well differentiated tissue observed in *PIK3CA*^*H1047R/H1047R*^ tumors was a focus of immature bone (B) in one. WT and *PIK3CA*^*WT/H1047R*^ tumors in contrast comprised variable admixtures of well-differentiated and organized tissue derivatives of all three germ layers. GI, gastrointestinal tissue; mAT, mouse adipose tissue (confirmed by independent mouse vs human immunostaining with Cyclophilin A, see Figure S7b); PE, pigmented epithelium; RE, respiratory epithelium; SBLs, sebaceous-like tissue. See also Figure S7 and Table S1. **b.** The Cancer Genome Atlas was used to extract genomic data from *PIK3CA*-associated cancers. These were analyzed in aggregate for the presence or absence of mutant *PIK3CA* alleles, followed by stratification of *PIK3CA* mutant-positive samples based on the presence of multiple mutant alleles, including cases where the wild-type (WT) *PIK3CA* allele is lost (WT-). Alternatively, *PIK3CA* mutant-positive samples were screened for multiple distinct *PIK3CA* mutations (*) or for the presence of additional mutations in proximal PI3K pathway components. **c.** Schematic of proximal class IA PI3K signaling.

## Discussion

We present the first pluripotent stem cell model permitting assessment of the consequences of selective genetic p110α activation specifically in a human developmental context. This is important given the well-documented differences between the pathways regulating mouse and human stem cell pluripotency and differentiation (51). By using CRISPR-mediated knock-in of *PIK3CA*^*H1047R*^ into one or both endogenous *PIK3CA* alleles, we were able to examine the importance of mutant *PIK3CA* allele dosage for pathway activation and downstream cellular responses. Human pluripotent stem cells are useful not only for study of human embryogenesis but also of the effects of pathological PI3K signaling, as seen in PROS and cancer cells (52). The model we have generated may thus be useful for understanding oncogenic actions of *PIK3CA*^*H1047R*^ in different contexts. By using expression from endogenous loci, by studying multiple clones of each genotype, and by controlling for non-specific variation introduced during the targeting process, we have minimized analytic problems arising from overexpression of the gene of interest and from non-specific genetic and chromosomal abnormalities.

*PIK3CA*^*H1047R*^ increased PI3K signaling “tone” both in growth factor-replete and growth factor-depleted medium. Most strikingly, we report distinct allele dose-dependent effects of mutant *PIK3CA* on stemness and pluripotency *in vitro* and *in vivo*, with a corresponding major alteration of the transcriptome triggered at a threshold between heterozygous and homozygous p110α activation. At odds with our finding in human stem cells, heterozygous expression of *PIK3CA*^*H1047R*^ in a human MCF10A breast epithelial cell line has previously been shown to cause widespread transcriptional changes, illustrating the notion that small changes in a non-linear system can have extensive consequences (53, 54). However, the mutant cells in these studies also had amplification of chromosome 5p13-15 (54), a region harboring the gene encoding the catalytic subunit of telomerase. This could have contributed to the observed discrepancy to our study. Alternatively, thresholds at which p110α signaling triggers its transcriptional effects may differ among cell types. Exemplifying this, either overexpression or endogenous expression of *PIK3CA*^*H1047R*^ induces multipotency in mammary tumors (8, 16), with the tumor cell of origin dictating phenotypic severity.

Although we describe the first stem cell-based study focusing on endogenous expression of the commonest pathogenic *PIK3CA* allele, several other studies have adopted different strategies to activate other components ofthe PI3K/AKT signaling cascade in this cell type (40, 55–58). Self-sustained stemness is a common motif in the phenotypes reported, and some studies, like ours, argue for discernible PI3K dose-dependency. For example, mouse pluripotent stem cells with homozygous knockout of the non-specific type IA PI3K negative regulator *Pten* exhibit impaired differentiation *in vitro* and *in vivo*, but this is not seen in heterozygous knockout cells (57). How strong PI3K activation sustains stemness remains to be determined, however our data suggest that induction of TGFβ/NODAL signaling is likely to be important. Supporting this, several transcriptional changes observed in *PIK3CA*^*H1047R/H1047R*^ cells were reciprocal to those in human pluripotent stem cells exposed to pharmacological inhibition of TGFβ signaling (59). It is also possible that the direct link between PI3K activation and *NODAL* expression underlies the previously reported association between PI3K/AKT activation and expression of *NANOG* (56, 60), a key pluripotency gene controlled by SMAD2/3 (61).

Our report of marked allele dose-dependent effects of *PIK3CA*^H1047R^ may have implications for understanding of PI3K-associated cancers. Many human cancers feature oncogenic alterations in *PIK3CA*, and not only are these frequently present in more than one copy, but they are also frequently accompanied by mutations of other pathway components, suggesting that cancer cells benefit from additional PI3K pathway activation. Future studies of the role of the PI3K pathway in cancer progression should incorporate consideration of PI3K signaling “dose” and the possibility of clear thresholds for biological consequences. Such considerations echo recent reports that an increased dosage of mutant *KRAS* influences clinical outcome and therapeutic targeting (62, 63). Based on our observations in human pluripotent stem cells with homozygosity for *PIK3CA*^*H1047R*^, it will be interesting to determine whether cancers with stronger activation of PI3K exhibit more aggressive features such as a higher degree of dedifferentiation and metastatic potential.

In contrast to the complex genetics of cancer, activating *PIK3CA* mutations arise heterozygously and in isolation in the severe overgrowth disorders known as PROS. An excess risk of adult cancer has not been reported in these mosaic disorders, in line with the notion that heterozygosity for *PIK3CA*^*H1047R*^ alone is not sufficient to cause cellular transformation. PROS also illustrates the importance of control of p110α signaling in early human development. Overgrowth in PROS commonly affects mesodermal and neuroectodermal lineages but rarely endoderm-derived tissues, prompting speculation that a sustained increase in PI3K activation impairs endoderm development (5). It has also been reported that class IA PI3K signaling is incompatible with directed definitive endoderm formation from human pluripotent stem cells, although this assertion is largely based on use of nonspecific pan-PI3K inhibitors (46, 47). In our study, we found no evidence that genetic PI3K activation impairs guided definitive endoderm formation in culture. Moreover, *PIK3CA*^*WT/H1047R*^ pluripotent stem cells gave rise to teratomas featuring well-differentiated endodermal components, arguing against a cell-autonomous defect in endoderm specification as an explanation for overall lack of endodermal overgrowth in PROS. The relatively mild biochemical and transcriptional consequences of heterozygous *PIK3CA* activation in stem cells, and their grossly normal early differentiation in several different experimental contexts suggests that any negative selection in certain lineages may be exerted only at later stages of differentiation. In contrast, homozygosity for *PIK3CA*^*H1047R*^ in early development will likely be selected against due to impaired differentiation.

In summary, this study demonstrates that the cellular consequences of the most common oncogenic *PIK3CA* mutation are allele dose-dependent. The observed near binary differences between *PIK3CA^H1047R^* heterozygosity and homozygosity suggest that cells may have a PI3K signaling threshold that determines the pathological consequences of this variant in development and cancer.

## Supporting information

## Acknowledgements

We thank Cornelia Gewert, James Warner from the Metabolic Research Laboratories (MRL) Histology Core and Gregory Strachan from the MRL Imaging Core. Technical assistance for this project was also provided by the UCL Cancer Institute Pathology Core Facility which is supported by the Cancer Research UK-University College London (CRUK-UCL) Centre Award [C416/A25145]. We thank Dr John Tadross and Dr Cheryl Scudamore for help with histopathological studies, Dr Marcella Ma and Dr Brian Lam from the MRL Genomics/Transcriptomics Core for RNA sequencing library preparation and raw read processing. We are grateful to Dr Adam Kashlak and Dr Matt Castle for advice on statistics and data processing, and to Dr Sarah Harrison, Dr Anne-Claire Guenantin and Dr Mo Mandegar for advice on stem cell maintenance and differentiation.

## Author contributions

Conceptualization: R.R.M., R.K.S. Methodology: R.R.M., R.K.S. Computation: R.R.M., S.P., N.M. Formal analysis: R.R.M., R.G.K., B.M.A., S.P., N.M. Investigation: R.R.M, R.G.K., W.P. Resources: B.V., R.K.S. Data curation: R.R.M., R.G.K., S.P., N.M. Manuscript preparation: R.R.M., R.K.S. Manuscript review/editing: R.R.M., B.V., R.K.S. Visualization/data presentation: R.R.M. Supervision: R.K.S. Project administration: R.R.M., R.K.S. Funding acquisition: R.R.M., B.V., R.K.S.

## Methods

### EXPERIMENTAL MODEL AND SUBJECT DETAILS

CRISPR/Cas9 targeting was performed on the male WTC11 induced pluripotent stem cell line (iPSC) line, a kind gift from Bruce Conklin (Gladstone Institute and UCSF). The derivation of this line has been described (64), and publicly available RNA, whole-exome and whole-genome sequencing data are available *via* the Conklin Lab’s website (https://labs.gladstone.org/conklin/pages/wtc-information) or *via* the Coriell Institute (cat. no. GM25256). In the current work, the parental line was used for gene editing at passage numbers P37 and P38. The derived iPSCs were used for experiments between P45 and P60.

The PROS patient-derived iPSC lines M98-WT and M98-E418K were obtained from a female, 18-year-old PROS patient by episomal reprogramming of a dermal fibroblast culture with 32 % mosaicism for *PIK3CA-* E418K. All clones used for experimental studies were confirmed transgene-free and expressed high levels of PSC-specific markers, comparable to those of a reference human pluripotent stem cell (hPSC) line. Karyotyping on a single line from each genotype confirmed lack of microscopic genetic rearrangements.

Cells were grown at 37 °C and 5 % CO_2_ in Essential 8 Flex (E8/F) medium on Geltrex-coated plates, in the absence of antibiotics. For maintenance, 70-90 % confluent cells were passaged as aggregates with ReLeSR, in the presence of the ROCK inhibitor RevitaCell (E8/F+R) during the first 24 h to promote survival. For experiments that required precise control of cell numbers, hPSC colonies were dissociated into single cells with StemPro Accutase, prior to manual cell counting.

All cell lines were tested negative for mycoplasma and genotyped routinely to rule out cross-contamination during prolonged culture. STR profiling was not performed.

### METHOD DETAILS

#### CRISPR/Cas9 targeting of hPSCs

hPSCs were targeted with plasmid-delivered wild-type Cas9 (pX459, Addgene #48139) and gBlock-encoded FE-modified sgRNAs (65). Targeting was performed by nucleofection of 5 μg pX459 plasmid (Cas9 wild-type), 3 μg sgRNA-encoding gBlock and either 200 pmol targeting template (for homozygous targeting) or a combination of 100 pmol targeting and “mock” templates (for heterozygous targeting). The nucleofected cells were seeded into Geltrex-coated 96-well plates and processed for sib-selection when ready for passaging. Sib-selection was performed as described previously (66), using 25-100 cells/well in each subcloning round. Wild-type iPSC lines obtained in the process of subcloning were banked as genetically-matched controls.

#### Embryoid body (EB) differentiation assays

EBs were established either by spontaneous self-aggregation of hPSCs or by forced aggregation into AggreWell plates. For self-aggregation, 50-70 % confluent hPSCs were dissociated into aggregates with ReLeSR, and the entire cell suspension from a 6-well transferred to one 60 mm Nunclon Sphera ultra-low attachment dish in Essential 6 (E6) medium supplemented with 0.4 % (w/v) polyvinylacohol (PVA) and RevitaCell (E6/PVA+R). EBs formed within 24 h, after which the medium was exchanged with E6 (without PVA and RevitaCell). The medium was exchanged again on day 3 of EB formation. For adherent outgrowths, the EBs were transferred to Geltrex-coated 6-well plates on day 4, either in regular E6 or in E6 supplemented with 10 % (v/v) fetal bovine serum (FBS), 100 nM BYL719 or 0.01 % (v/v) DMSO. The EBs from a single Nunclon Sphera dish were used to seed four wells of a 6-well plate or eight wells of a 12-well plate. EB outgrowths were collected for RNA extraction on day 10 of EB formation. In one experiment, suspension EBs were also collected on day 4 and day 13.

EB set-up in AggreWell plates followed the manufacturer’s instructions, with E6/PVA+R as medium for cell seeding. A total of 2.4 × 10^5^ cells were seeded in each well, for a final density of 200 cells/microwell. EBs formed within 24 h, and the contents of four or five individual wells were transferred to a single Nunclon Sphera ultra-low attachment dish for culturing in either mesoderm (10 ng/ml BMP4, 5 ng/ml Activin A, 5 ng/ml FGF2) or endoderm (0.25 ng/ml, 100 ng/ml Activin A, 2.5 ng/ml FGF2) induction medium. After three days of induction, the EBs were transferred to Geltrex-coated 6-well plates for adherent growth and maintained in E6 until day 10. Cells were collected for RNA extraction on day 0 (iPSC stage), day 4, day 7 and day 10 of EB formation.

For immunocytochemistry (see below), day 4 EBs were also seeded for adherent growth in Geltrex-coated 4-well or 35 mm Ibidi imaging and processed for staining on day 10.

#### Definitive endoderm differentiation

Definitive endoderm differentiation of hPSCs was carried out according to a modified version of the protocol described by (45). Cells were seeded in Geltrex-coated 12-well plates at densities between 5,000-7,500 cells/cm^2^ (WT, *PIK3CA*^*WT/H1047R*^, *PIK3CA*^*H1047R/H1047R*^) or 8,500 cells/well^2^ (WT, *PIK3CA*^*WT/E418K*^), seeding a minimum of two cultures per clone. Two days post-seeding, the cells were induced to differentiate in the presence of BYL719 (100 nM) or the corresponding dilution of DMSO. Samples were collected for RNA extraction at baseline (immediately before induction) and on each one of the following days of the differentiation protocol. In one experiment, cells were also fixed for immunofluorescence at the end of differentiation.

#### Tumor xenografts assays

Tumor xenografts were generated from a total of 10 iPSC cultures (*N* = 5 WT, *N* = 3 *PIK3CA*^*WT/H1047R*^, *N* = 2 *PIK3CA*^*H1047R/H1047R*^) by subcutaneous injection into immunodeficient, male NSG mice (Jackson #005557) at 12 weeks of age. Cells had been cultured according to standard procedures in Geltrex-coated T75 flasks, ensuring 95-100 % confluence on the day of injection. The cells were processed for aggregate dissociation with ReLeSR, collected in E8/F+R and centrifuged at 200g for 3 min. Each cell pellet was resuspended in 130 μl ice-cold E8/F+R, followed by mixing with 70 μl ice-cold human ESC-qualified Matrigel. From this suspension, 200 μl were used for injections within 30 min of preparation (kept on ice throughout), using pre-chilled syringes (20.5 gauge needles). Individual animals were culled when tumors reached approximately 1.4 cm^3^ in size, or if they became ill suddenly. All animal procedures were performed with approval from the local Animal Welfare Ethical Review Body (AWERB), and in accordance with Home Office regulations (The Animal [Scientific Procedures] Act 1986).

#### Tumor histopathology

Each tumor was processed for formalin fixation, paraffin embedding, microtome sectioning and hematoxylin and eosin (H&E) staining as described in (68). Individual tumors were cut in half before side-by-side paraffin embedding of the two halves, with the cut surfaces facing out to allow different areas of each tumor to be processed at the same time. A total of 12 sections (3 μm each) were made per paraffin block and mounted on Superfrost Plus slides, yielding a total of 24 different tumor areas because of side-by-side embedding. Odd-number slides were processed for H&E staining and the rest used for immunohistochemistry. The slides were analyzed blindly by a human pathologist and processed for automated brightfield imaging on an AxioScan Z1 (Zeiss) slide scanner.

#### RNA sequencing

RNA was extracted as described above, followed by quantification and quality assessment on an Agilent Bioanalyzer using the RNA 6000 Nano Kit according to the manufacturer’s instructions, confirming that all samples had a RIN score of 10. A total of 1 μg per sample was used to synthesize 50 bp long single end mRNA libraries with an Illumina TruSeq Stranded mRNA Library Prep Kit. The integrity and quantity of the libraries were determined on the Bioanalyzer using the DNA 12000 Kit. The barcoded libraries were pooled and sequenced on an Illumina HiSeq 4000, with an average depth of 20 million reads per sample. The raw reads were mapped to the human genome build GRCh38, and gene level counts were performed using Spliced Transcripts Alignment to a Reference (STAR) v2.5 (71). Subsequent data processing followed the method outlined in (72).

#### Pathway analyses

Ingenuity^®^ Pathway Analysis (build version 448560M, content version 36601845) was conducted on the 1,914 differentially expressed genes in *PIK3CA*^*H1047R/H1047R*^ iPSCs, using the Ingenuity Knowledge Base (Genes Only) as reference set and including both direct and indirect relationships. Relationships were only considered if experimentally observed. The results from the Upstream Regulator Analysis (73) were exported and processed for visualization in RStudio. Independently, the same set of genes were used for pathway analysis with the CytoScape plug-in ClueGO (v2.3.4), focusing on pathways within the KEGG ontology (Build 01.03.2017) and applying the following parameters: merge redundant groups with 50 % overlap; evidence codes used: all; statistical test: enrichment/depletion (two-sided hypergeometric test) with Benjamini-Hochberg correction; minimum number of genes per term = 10; minimum percentage enrichment = 4; Kappa Score Threshold = 0.4. The created network represents the pathway terms as nodes linked based on their term-term similarity, as determined by the Kappa Score. The size of the nodes reflects the enrichment significance of the terms, and only terms with FDR ≤ 0.05 are shown.

### TCGA data analysis

Somatic mutation tables (MAFs) from whole-exome sequencing data across 11 cancer types (BLCA, BRCA, CESC, CRC, ESCA, GMB, HNSC, LUSC, STAD, UCEC, UCS) were downloaded from the TCGA portal through the Genomic Data Commons (GDC) Data Transfer Tool. Mutation calls generated by Varscan2 (74) were used. To limit false positives, for those variants with a VAF (t_alt_count/t_depth) < 0.05, we retained those that were also identified by the MuTect2 algorithm (75). Functional annotation of genomic variants was performed with ANNOVAR (76). Purity, ploidy and copy number profiles of tumor cells were obtained with ASCAT (77) run using default parameters on SNP6.0 data.

Mutation multiplicity, or mutation-copy-number, describing the number of mutant copies of a mutation, is a function of the purity of the sample, the ploidy of the tumor cells and the relative frequency of the mutation (i.e. the VAF). The mutation multiplicity was calculated as previously described (78), using the following equation:

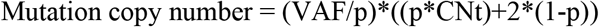

Where CNt is the local copy number at the mutated base and p is the estimated purity of the tumor sample. Mutations exhibiting a mutation multiplicity ≥ 1.5 were classified as ‘gained’. If the mutation multiplicity was equal the local copy number and the genomic segment harboring the mutation was subject to LOH, the wildtype allele was inferred to be absent.

### DATA PROCESSING

Data processing and visualization were performed in RStudio (https://www.rstudio.com/). Individual measurements, rather than summary statistics, are displayed in all cases except for the RNAseq data. All raw data and bespoke RNotebooks containing guided scripts used to analyze larger datasets are available *via* the Open Science Framework (link to be added upon acceptance). The original RNAseq data have been deposited in the GEO, under accession number: GSEXXXXXX (to be done upon acceptance). All uncropped Western blots are provided as a separate supplemental file if required. This includes both blots that are displayed in the paper as well as additional replicates.

## Supplemental Material

Separate PDF file with supplemental figures and tables

Separate PDF file with supplemental methods

Separate PDF file containing key resource information, including catalogue numbers, is available for upload upon acceptance.

Separate Word file containing all uncropped Western blots, including additional replicates, is available for upload upon acceptance

