## Supplementary material for "Oncogenic *PIK3CA* promotes cellular stemness in an allele dose-dependent manner"

### Supplemental Figures

a

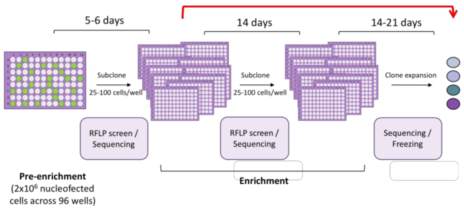


b

| Off-target site (prediction algorithm) | Off-target sequence (sgRNA-Fb: ATGAATGATGCACATCATGG) |
|---|---|
| Off-target 1: RP11-796E10.1 (Cas-OFFinder) | ATGAATGtAaGCACATaATGG-NGG |
| Off-target 2: *ACADVL* (Cas-OFFinder) | ATGAATGCATcCAaATCATGG-NGG |
| Off-target 3: *NEURL4* (Cas-OFFinder) | ATGACAcGATGCACATCcTGG-NGG |
| Off-target 4: *KALRN* (Cas-OFFinder) | ATGAATGATGAACACATagTGG-NGG |
| Off-target 5: RP11-25H12.1 (Cas-OFFinder) | ATGAATGATGCCCACATgATtG-NGG |
| Off-target 6: *ADAM19* (Cas-OFFinder) | gTGAATGATaCACATTTCATGG-NGG |
| Off-target 7: *MGRN1* (Cas-OFFinder) | ATGAACATGgTGCACATCtTGG-NGG |
| Off-target 8: *LHFP* (Cas-OFFinder) | ATGAATGtTGCcAGCATCATGG-NGG |
| Off-target 9: *ANO4* (Cas-OFFinder) | ATGcATcATcCACATCATGG-NGG |
| Off-target 10: *IQCJ-SCHIP1* (Cas-OFFinder) | AgGAATGATGCAaAgCATGG-NGG |
| Off-target 11: *SEPT7* (Cas-OFFinder) | ATGAtTGATGCACATCATtt-NGG |
| Off-target 12: *TBCK* (Cas-OFFinder) | ATGAATGATGaACATCAacG-NGG |
| Off-target 13: *ELF2* (Cas-OFFinder) | ATGAAgGATGCACATCtgGG-NGG |
| Off-target 14: *RP11-122F24.1* (Cas-OFFinder) | ATGAATtATGtACATCATtG-NGG |
| Off-target 15: *CENPP* (Cas-OFFinder) | ATGAATGAaGCAgATCATGa-NGG |
| Off-target 16: *ARMC10* (crispr.mit.edu) | AgGCATGtTtCACATCATGG-NAG |
| Off-target 17: *GRM1* (crispr.mit.edu) | AatAATGATGtACATCtTGG-NAG |
| Off-target 18: *TTC21B-AS1* (crispr.mit.edu) | gTGAATGtgGCACATCATtG-NAG |

c

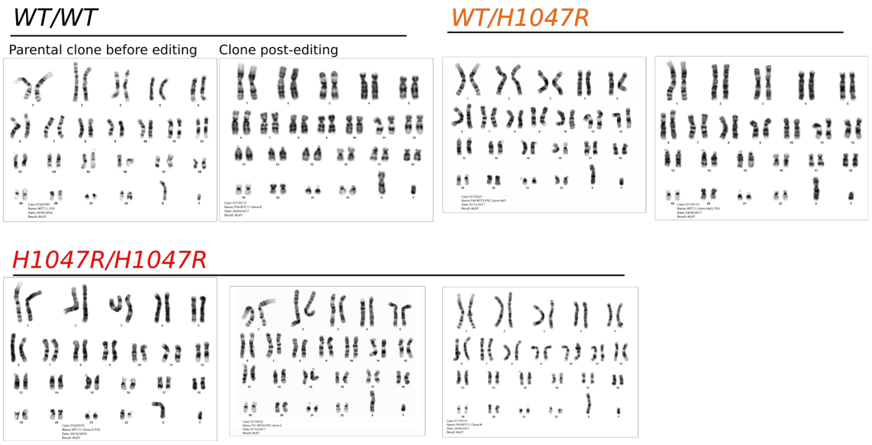


**Figure S1, related to Figure 1. CRISPR editing in human iPSCs to knock-in *PIK3CA*$^{\mathbf{H1047R}}$. a.** Schematic of the sib-selection protocol used to isolate correctly targeted human iPSCs. In some cases, clonal lines were obtained without a need for a second round of subcloning (red arrow). **b.** A table of the 18 computationally predicted off-target sites that were examined in 2 wild-type, 3 *PIK3CA*$^{WT/H1047R}$ and 6 *PIK3CA*$^{H1047R/H1047R}$ clones. The following off-target sites were located in coding regions: 1, 2, 3, 16, 17 and 18. The sequencing failed for *SEPT7*, Off-target 11, due to poor primers. A heterozygous SNP (dbSNP ID: rs76736990) in *KALNR*, Off-target 4, was observed in all clones, suggesting that it was present in the original donor and not a consequence of CRISPR targeting. This was confirmed by examining the publicly available whole-genome sequencing data for the parental WTC11 cell line. **c.** Metaphase spreads confirming a normal 46XY karyotype in wild-type and *PIK3CA*$^{H1047R}$ iPSC clones following gene editing. A total of 20 metaphases were examined in all cases except the parental clone before editing (11 metaphases analyzed).

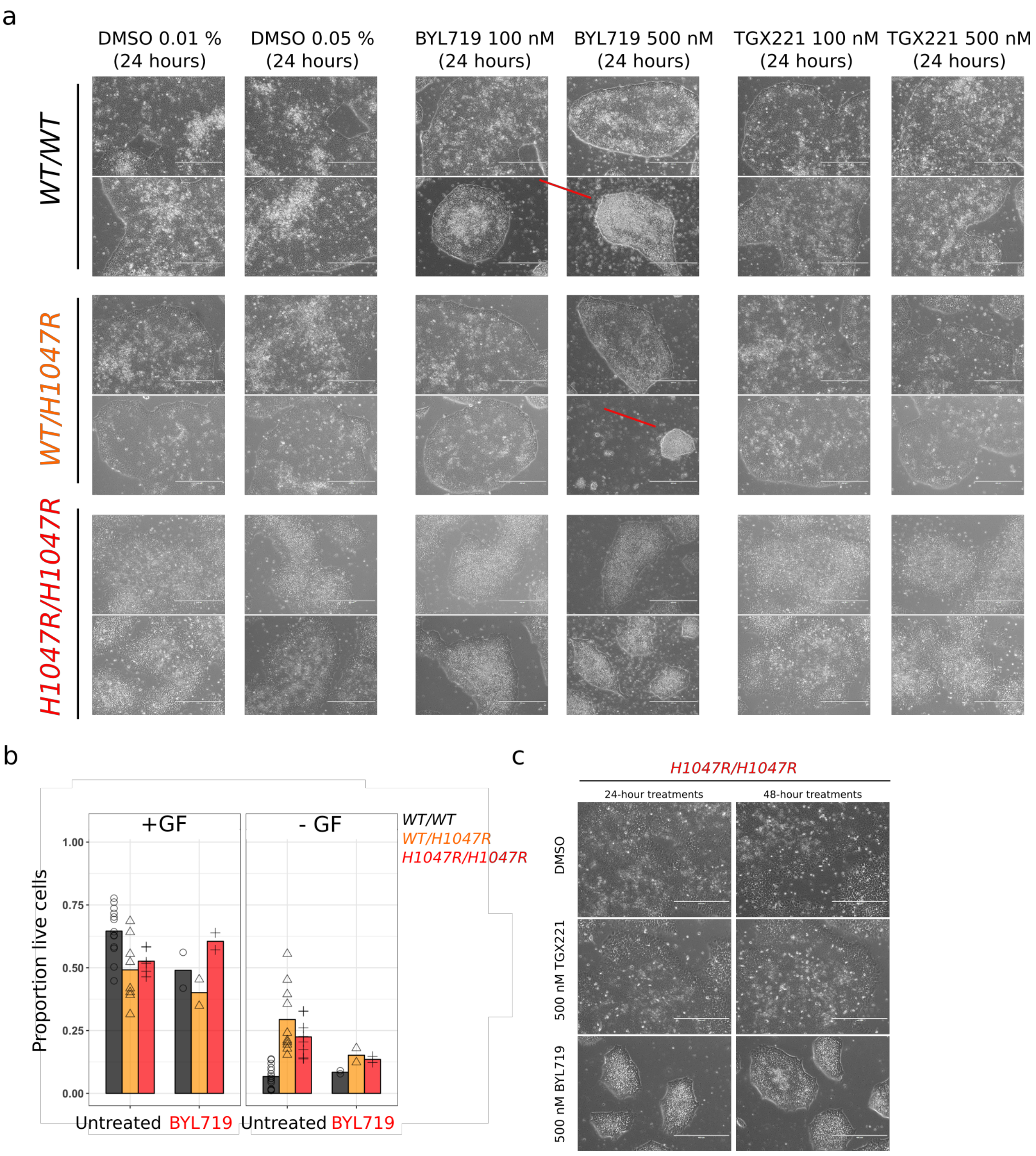


**Figure S2, related to Figure 2. Drug response and survival phenotypes of wild-type and *PIK3CA*$^{\mathbf{H1047R}}$ iPSCs. a**. The morphological appearance of the clones used to obtain the signaling data in Figure 2c. Higher concentrations of BYL719 (500 nM) had cytotoxic effects in some of the wild-type (*WT/WT*) and *PIK3CA*$^{WT/H1047R}$ (*WT/H1047R*) clones (red arrows). This was not observed for *PIK3CA*$^{H1047R/H1047R}$ (*H1047R/H1047R*) clones. **b**. Cell viability at baseline and in response to 24-h growth factor (GF) depletion, +/- BYL719 (100 nM), was quantified by Annexin V staining of apoptotic cells combined with cell size discrimination of dead cells. Except for BYL719 treatments, which were performed in one experiment using technical duplicates of a single clone per genotype, data for the remaining treatments are from 3 independent experiments and at least 3 iPSC clones per genotype. Data points correspond to individual cell cultures, with bars representing the mean percentage of live cells for a given condition. **c**. The morphological appearance of *PIK3CA*$^{H1047R/H1047R}$ (*H1047R/H1047R*) clones treated with DMSO (control), 500 nM TGX221 (p110β inhibitor) or 500 nM BYL719 (p110α inhibitor). Representative of at least 3 experimental replicates and 6 *PIK3CA*$^{H1047R/H1047R}$ clones.

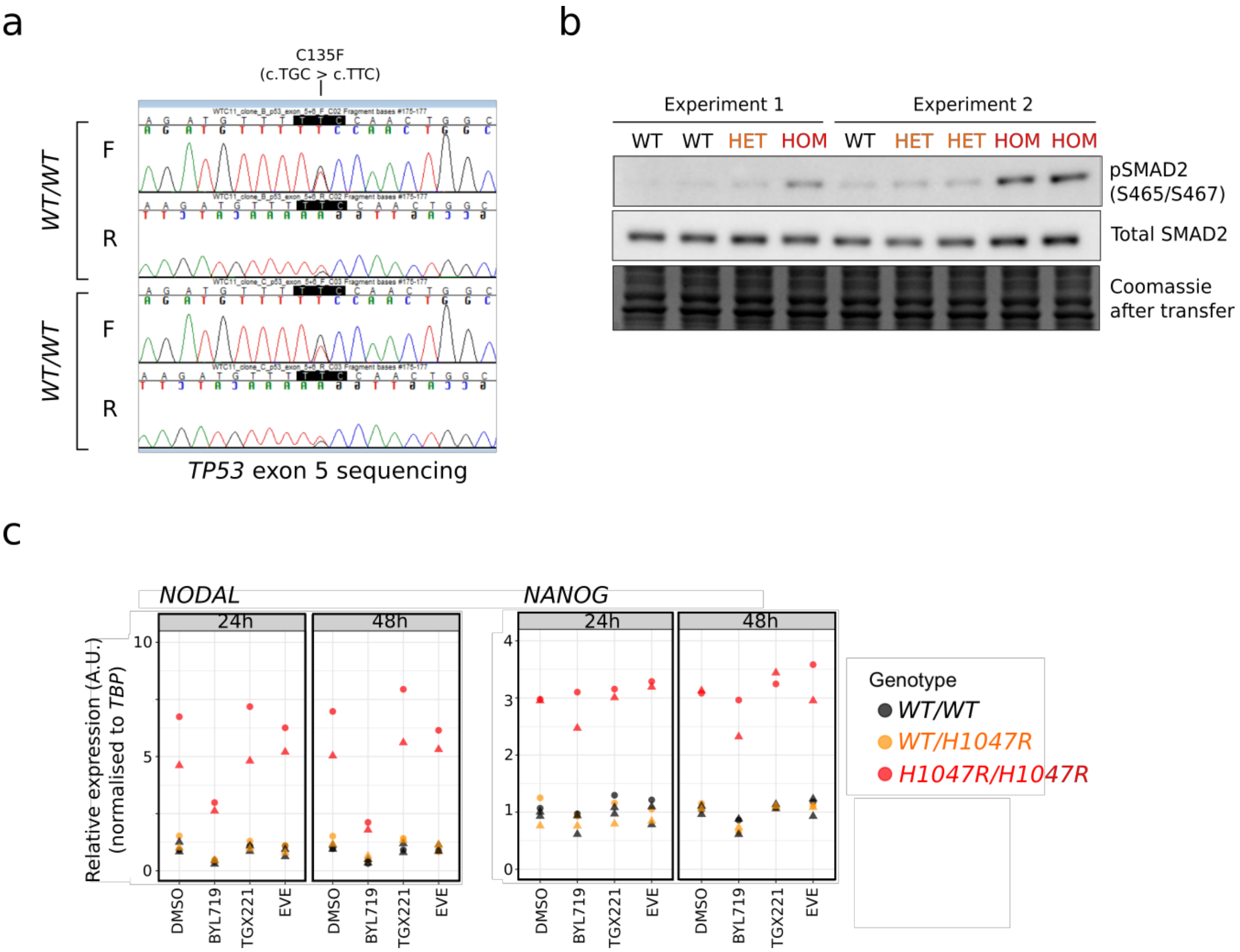


**Figure S3, related to Figure 3. *TP53* sequencing, SMAD2 phosphorylation and regulation of *NODAL* and *NANOG* gene expression in wild-type and *PIK3CA*$^{H1047R}$ iPSCs. a.** Sanger sequencing traces of *TP53* exon 5 in two of the *PIK3CA* WT cultures that had acquired a heterozygous *TP53*-C135F variant in culture. F, forward primer. R, reverse primer. **b**. Western blots for pSMAD2 (S465/S467) and total SMAD2 in wild-type (WT), *PIK3CA*$^{WT/H1047R}$ (HET) and *PIK3CA*$^{H1047R/H1047R}$ (HOM) iPSCs. Two experimental replicates are shown, conducted on 2 wild-type, 3 *PIK3CA*$^{WT/H1047R}$ and 2 *PIK3CA*$^{H1047R/H1047R}$ clones. **c**. RT-qPCR analysis of *NODAL* and *NANOG* expression in wild-type (WT), *PIK3CA*$^{WT/H1047R}$ and *PIK3CA*$^{H1047R/H1047R}$ iPSCs following 24 h and 48 h treatments with either DMSO (control), 500 nM BYL719 (p110α inhibitor), 500 nM TGX221 (p110β inhibitor) or 5 nM everolimus (mTORC1 inhibitor). The data are from 2 independent experimental replicates, with 2 wild-type, 1 *PIK3CA*$^{WT/H1047R}$ and 2 *PIK3CA*$^{H1047R/H1047R}$ clone(s).

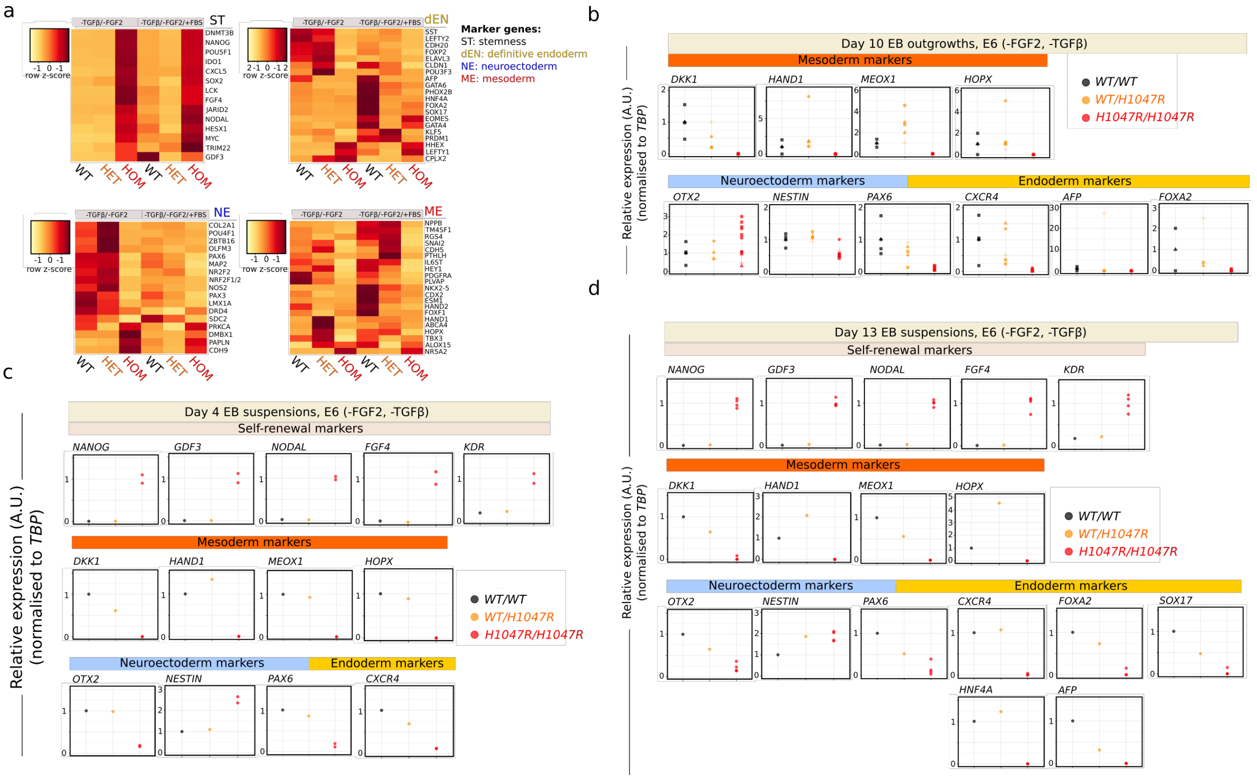


**Figure S4, related to Figure 4. Lineage-specific gene expression in self-aggregating wild-type or *PIK3CA*$^{H1047R}$ EBs under different conditions and at different time points. a**. Real time quantitative PCR scorecard-based profiling of lineage markers in wild-type (WT), *PIK3CA*$^{WT/H1047R}$ (HET) and *PIK3CA*$^{H1047R/H1047R}$ (HOM) EB outgrowths collected on day 10 of EB formation. The EBs were maintained in E6 throughout the protocol or in E6 supplemented with 10 % fetal bovine serum (FBS) from day 4 of EB formation, at the onset of adherent outgrowth formation. Gene heatmaps are shown across rows grouped according to lineage. Colors correspond to expression z-scores as indicated. The data are from a single experiment with one clone per genotype. **b**. As in Figure 5d but assessing additional lineage-specific markers as indicated. The data are from day 10 EB outgrowths maintained in E6 throughout the experiment. **c**. RT-qPCR analysis of the indicated lineage-specific markers in day 4 suspension EBs maintained in E6. The data are from a single experiment with one wild-type (WT), one *PIK3CA*$^{WT/H1047R}$ and 2 *PIK3CA*$^{H1047R/H1047R}$ clones. **d.** RT-qPCR analysis of lineage-specific markers in day 13 suspension EBs maintained in E6. The data are from one experiment with one wild-type (WT), one *PIK3CA*$^{WT/H1047R}$ and 4 *PIK3CA*$^{H1047R/H1047R}$ clones. Endoderm formation occurred at later stages of spontaneous EB differentiation, and the number of endoderm makers in each panel reflects the ability to detect these genes above background at each time point. All RT-qPCR expression values are normalized to *TBP* and scaled internally. Expression values are in arbitrary units (A.U.).

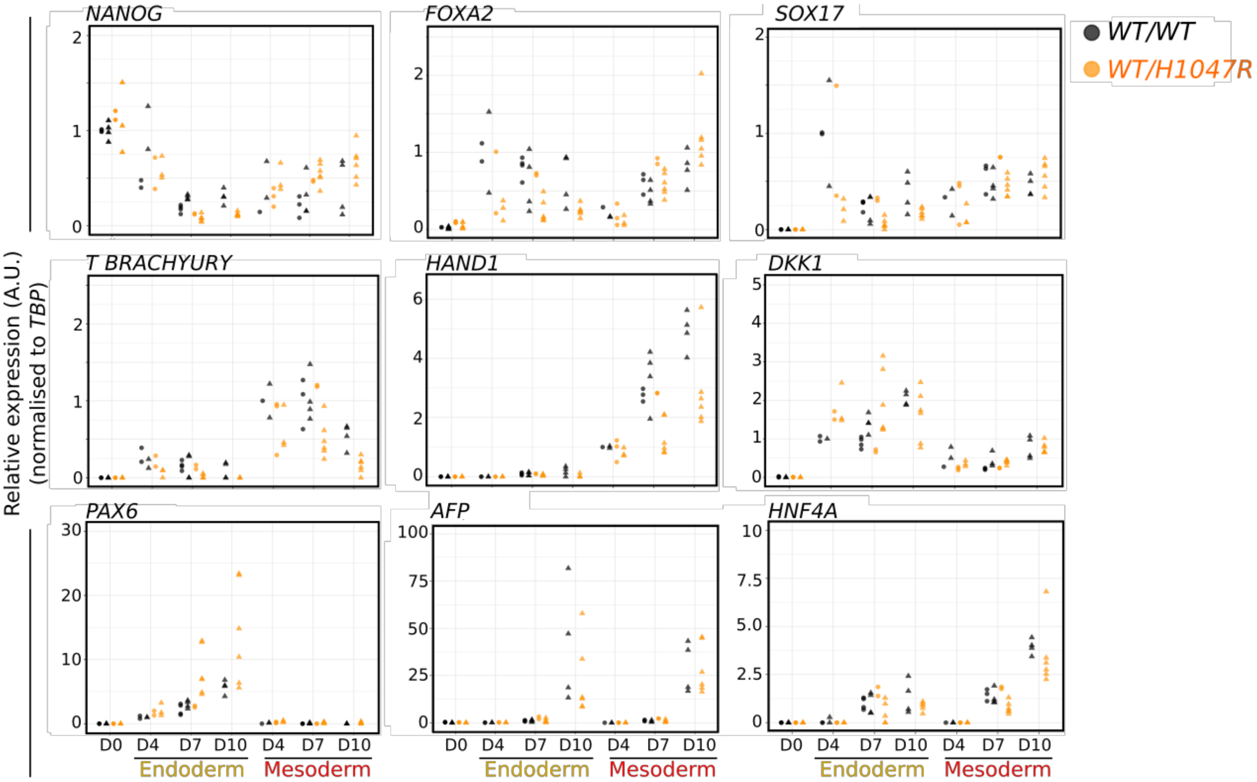


**Figure S5, related to Figure 5. Lineage-specific gene expression in endo- or mesoderm-induced wild-type or *PIK3CA*$^{WT/H1047R}$ EBs.** Real time quantitative PCR assays of samples used in Figure **5b** plus additional replicates and time-points. Data shown for each timepoint represent 4-8 replicates from 2-4 clones. Gene expression values were normalized to *TBP* and scaled internally. The final values are in arbitrary units (A.U.). D, day.

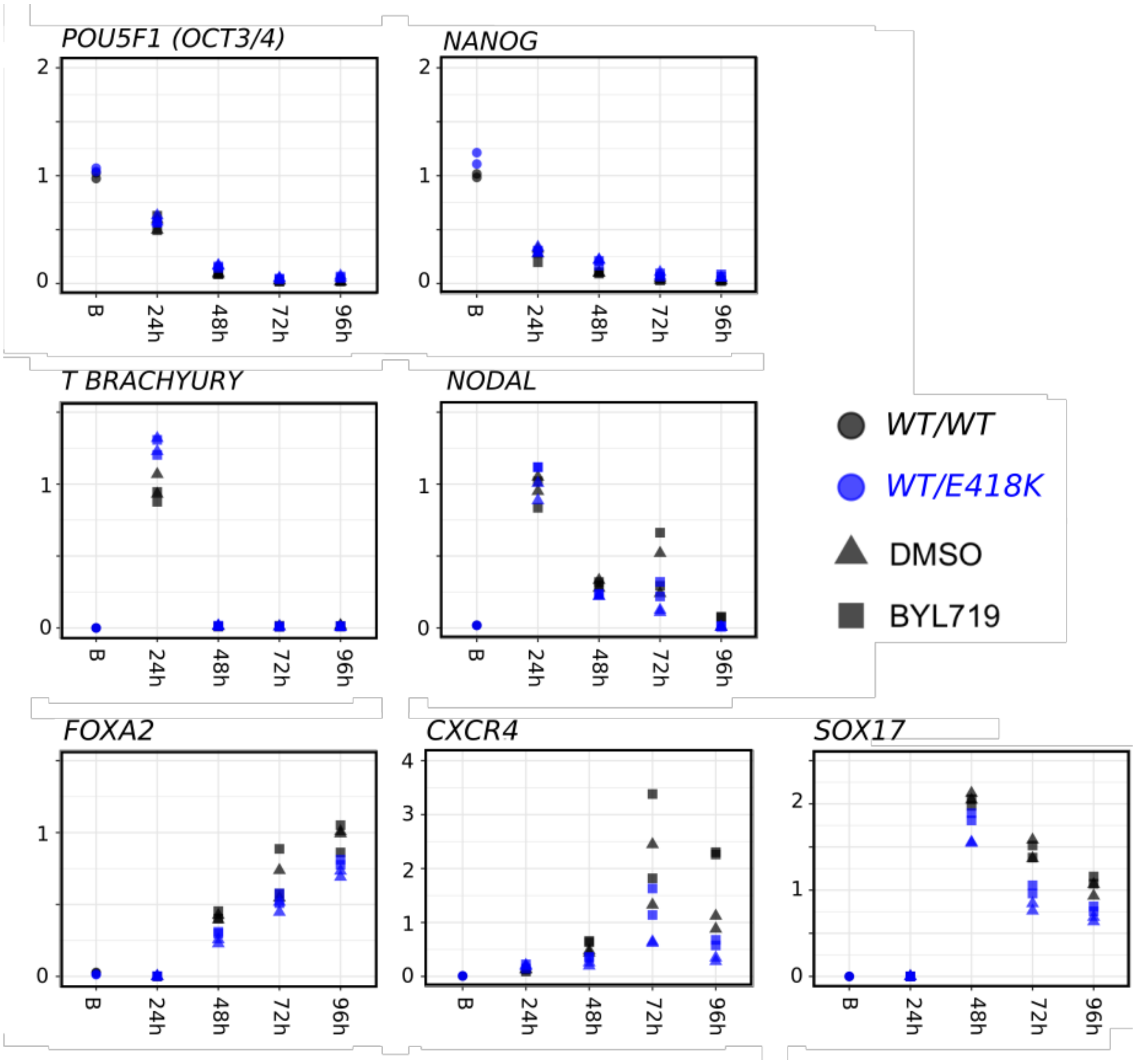


**Figure S6, related to Figure 6. Gene expression analysis of monolayer-based endoderm differentiation of isogenic patient-derived iPSCs (wild-type or *PIK3CA*$^{WT/E418K}$).** RT-qPCR gene expression analysis of lineage-specific markers throughout the course of monolayer-based definitive endoderm formation in genetically-matched WT or *PIK3CA*$^{WT/E418K}$ iPSCs derived from a female PROS patient. Differentiations were performed in the presence of DMSO (control) or 100 nM BYL719. The data are from a single experiment with one clone per genotype, with duplicate cultures of each clone.

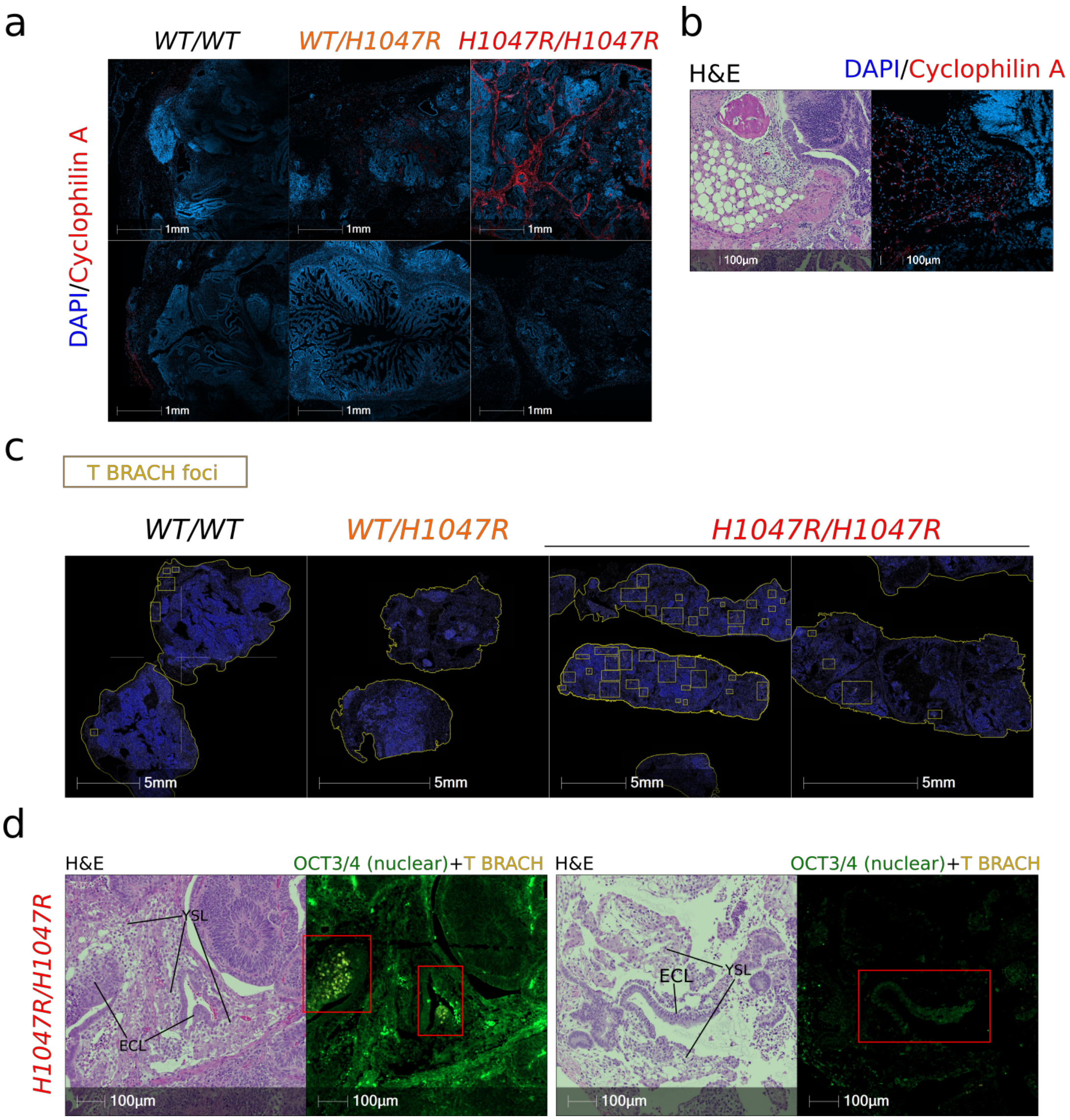


**Figure S7, related for Figure 7. Characterization of xenograft tumors derived from wild-type or *PIK3CA*$^{H1047R}$ iPSCs. a**. Cyclophilin A-based detection of mouse host tissue in immunofluorescent-stained slides from human iPSC-derived tumor xenograft. DAPI was used for nuclear visualization. The immunofluorescence micrographs are representative of 4 WT (*WT/WT*) tumors from 3 different iPSC clones, 3 *PIK3CA*$^{WT/H1047R}$ (*WT/H1047R*) tumors from 3 different clones, and 2 *PIK3CA*$^{H1047R/H1047R}$ (*H1047R/H1047R*) from 2 different clones. Scale bar = 1 mm. **b.** Example of matched hematoxylin and eosin (H&E) and DAPI/Cyclophilin A-stained slides from a wild-type tumor, demonstrating that the observed adipose tissue is of mouse origin. Representative of observations in the remaining tumors. **c**. Automated counting of T BRACHYURY (T BRACH)-positive cells in immunofluorescent stained slides from human iPSC-derived tumor xenografts. Areas with increased density of T BRACH-positive cells were marked manually as indicated by the yellow rectangles. All tumor slides are shown. Scale bar = 5 mm. **d**. Matched H&E and OCT3/4/T BRACHYURY immunofluorescent stained slides from the two *PIK3CA*$^{H1047R/H1047R}$ tumors. The OCT3/4 signal can be distinguished by its nuclear localization. Scale bar = 100 µm. ECL, embryonal carcinoma-like tissue; YSL, yolk sac-like tissue.

| | Endoderm | | | Mesoderm | | Ectoderm | | | Other | | Overall maturity |
|---|---|---|---|---|---|---|---|---|---|---|---|
| | GI epithelium | Respiratory epithelium | Immature unilayered tubular structures[a] | Cartilage | Bone | Pigmented epithelium [b] | Sebaceous-like glands | Neurones (ganglion cells) | Yolk sac-like elements | Necrosis [b] | |
| WT | | | + | + | + | 10 % | + | | | < 5 % | ++ |
| | | | + | | + | 15 % | | | | < 5 % | ++/+++ |
| | | | + | + | + | 10 % | + | + | | < 5 % | ++ |
| | | | + | | + | < 5 % | | | Focal (?) | < 5 % | +/++ |
| | | | + | | + | < 5 % | | | Focal (?) | < 5 % | +/++ |
| HET | + | + | + | + | | < 5 % | + | + | | < 5 % | +++ |
| | | + | + | + | Focal | 10 % | + | + | | < 5 % | ++ |
| | | + | + | | | 10 % | | + | | 10 % | ++ |
| HOM | | | + | | Focal | | | | Focal | 30 % | + |
| | | | + | | | | | | ++ | 20 % | -/+ |

**Table S1, Related to Figure 7 and Figure S7. Summary of the histopathological evaluation of wild-type and *PIK3CA*$^{H1047R}$ tumour xenografts.** The tumours were obtained from 5 wild-type (WT), 3 *PIK3CA*$^{WT/H1047R}$ (HET) and 2 *PIK3CA*$^{H1047R/H1047R}$ (HOM) iPSC cultures, using independent mutant clones and four wild-type clones. Six haematoxylin and eosin-stained slides with two different areas from each tumour were analysed blindly by a human pathologist. [a] Presumed endoderm. [b] The quantification was performed subjectively by the pathologist, and the numbers are only approximate. (?) Sparse and therefore less certain appearance. In addition to the indicated tissues, all tumours also contained mature adipose tissue and blood vessels. However, mouse-specific Cyclophilin A-staining showed that these were not of human origin (Figure S7b).
