## Supplementary material for "Oncogenic *PIK3CA* promotes cellular stemness in an allele dose-dependent manner"

### Supplemental methods

#### Karyotyping

For karyotyping, subconfluent hPSCs from two 6-wells were treated with 0.1 µg/ml KaryoMAX Colcemid solution for 90 min, dissociated into single cells, and the final cell pellet resuspended in hypotonic KCl solution (0.055 M) for 5 min at 37 °C. Next, the suspension was centrifuged at 200g for 5 min, and the pellet resuspended in freshly prepared cold fixative (3:1 methanol:glacial acetic acid). The fixed cell suspension was used for slide preparation and banding analysis by the East Anglian Medical Genetics Service.

#### Genotyping

For routine, confirmatory genotyping and for rapid post-gene editing mutation screening, genomic DNA (gDNA) from cells was extracted using QuickExtract solution according to the manufacturer's instructions. For high-quality sequencing, including screening for off-target and *TP53* mutations, gDNA was extracted with Qiagen's QIAamp DNA Micro Kit according to the manufacturer's instructions. The targeted DNA region was amplified by PCR with GoTaq Green according to the manufacturer's protocol, and the amplicon sequenced in both directions using the BigDye Terminator v3.1 Cycle Sequencing Kit following the manufacturer's protocol. For RFLP, the GoTaq Green PCR reaction was performed with a FAM-labelled version of the reverse primer. From the 25 µl PCR reaction, 10 µl were taken forward to a 30 µl digestion reaction with BmgBI, performed in line with the manufacturer's protocol. Next, 2 µl of each digestion reaction were mixed with 13 µl GeneScan 600 LIZ size standard stock, followed by capillary electrophoresis and fluorescent detection on an ABI 3730 sequencer (Thermo Fisher Scientific). The standard stock was prepared by mixing 1 ml HiDi formamide with 45 µl LIZ standards. Following electrophoresis, the results were extracted using the GeneMapper 5.0 software. The percentage of H1047R-positive cells was determined from individual peak areas with the following formula, with target peak corresponding to the signal from the BmgBI digestion product: E = (area of target peak) / [(area of target peak) + (area of unmodified peak)].

#### Off-target screening

Off-target sites were analyzed with the Zhang Lab's online sgRNA tool (October 2015). With the development of more advanced algorithms, the off-target list was re-evaluated in September 2017 using the Cas-OFFinder algorithm (67), selecting NGG as PAM site and allowing for up to five mismatches and two bulges against the human genome build hg38. The resulting list was filtered for targets with up to three mismatches, including up to two mismatches with two bulges. The top 15 off-target sites (both coding and non-coding) predicted by Cas-OFFfinder and the three coding off-target sites (among all top 50 off-targets) obtained with the Zhang Lab's tool were selected for confirmatory genotyping by Sanger sequencing in three WT clones, three $PIK3CA^{WT/H1047R}$ clones and six $PIK3CA^{H1047R/H1047R}$ clones. Additionally, a tiled primer matrix was designed, spanning 851 bp and 1126 bp upstream and downstream, respectively, of *PIK3CA* codon 1047. This was used to test for larger DNA rearrangements that could arise from CRISPR targeting. Arguing against such rearrangements, different primer combinations gave rise to an amplification product of the correct size in the profiled WT ($N = 1$), $PIK3CA^{WT/H1047R}$ ($N = 3$) and $PIK3CA^{H1047R/H1047R}$ ($N = 10$) clones. In addition, all PCR products from the WT clone and two $PIK3CA^{H1047R/H1047R}$ clones were Sanger sequenced, confirming lack of unintended indels.

As an additional check that the observed homozygosity for *PIK3CA* H1047R was not a false positive caused by loss of heterozygosity in the targeted cells, gDNA from WT and $PIK3CA^{H1047R/H1047R}$ iPSCs were mixed 1:1 and sequenced across *PIK3CA* codon 1047. This resulted in equally-sized chromatogram peaks for the WT and *PIK3CA* H1047R allele, consistent with homozygosity in the mutant cell lines. This check was performed with 7 $PIK3CA^{H1047R/H1047R}$ clones, mixed with either one of two independent WT clones.

#### Immunohistochemistry (IHC)

Formalin-fixed and paraffin-embedded teratoma sections were processed for dewaxing and rehydration, followed by high-pH antigen retrieval in DAKO Antigen Retrieval Solution (pH 9) in a water bath at 97 °C for 20 min. For Cyclophilin A-based detection of mouse host tissue (69), DAKO's Antigen Retrieval Solution (pH = 6) was used instead. Next, the slides in antigen retrieval solution were cooled down for 15 min at -20 °C, followed by a wash in Tris-buffered saline (TBS; 20 mM Trizma base, 150 mM NaCl, pH = 7.6) supplemented with 0.1 % Tween-20 (TBS/T) and two washes in DAKO wash buffer (5 min per wash). Slides were blocked in 2.5 % ready-to-use (R.T.U.) horse serum (HS) for 1 h at room temperature, using 300 µl blocking solution per slide. Primary antibody dilutions, including the corresponding isotype controls, were prepared in 2.5 % R.T.U. HS, and 200 µl were applied to each slide for overnight incubations at 4 °C. For co-staining, primary antibodies were applied simultaneously. The following day, the slides were placed at room temperature for 2 h, then washed five times in DAKO wash buffer, before room temperature incubation with the appropriate combination of secondary antibodies diluted 1:1,000 in 2.5 % R.T.U. HS. For dual staining requiring detection with goat anti-mouse and donkey anti-goat secondary antibodies, each slide was firsts incubated in donkey anti-goat antibody for 2h, followed by five washes in DAKO wash buffer, and another 2-h incubation in goat anti-mouse antibody, thus avoiding cross-reactivity between the two secondary antibodies. The second incubation was omitted when working with a single secondary antibody. After the final incubation, the slides were washed another five times and processed for overnight mounting in Fluoroshield DAPI, carried out at room temperature. The slides were processed for automated image acquisition on the AxioScan Z1 slide scanner, and the HALO software (Indica Labs) was used for contrast/brightness adjustments and visualization. The same adjustments were applied to all images obtained in the same staining round. The specificity of each signal was determined based on its subcellular localization and a comparison to the corresponding IgG control staining. Automated cell counting of T BRACHYURY-positive cell was conducted in HALO, following nuclear segmentation based on the DAPI signal. The analysis was performed on entire tissue slides.

**Immunocytochemistry (ICC)**

Adherent cells were fixed for 15 min with 4 % (v/v) formaldehyde, followed by permeabilization in PBS supplemented with 0.1 % (v/v) TritonX-100 (PBS/T) for 15 min and 60-90 min blocking in PBS/T with 5 % (v/v) horse serum (HS; PAN-Biotech). Primary antibodies were diluted in PBS with 1 % (v/v) HS and used in overnight incubations at 4 °C. For co-staining, multiple primary antibodies were applied simultaneously. The following day, the samples were equilibrated to room temperature, washed three times in PBS and incubated with the appropriate fluorophore-conjugated secondary antibodies diluted in PBS with 1 % HS, for 2 h away from light. Co-staining with secondary antibodies with the potential to cross-react was performed as outlined in the previous section, with PBS used as wash buffer and PBS with 1 % HS as dilution buffer. Anti-rabbit and anti-mouse secondary antibodies were diluted 1:1,000, and the anti-goat antibody 1:2,000. Next, cells were incubated in CytoPainter Phalloidin-iFlour 555 (1:1,000) for 45 min for F-Actin visualization, followed by nuclear counterstaining with Hoechst, DAPI (Nuclear Select FX Labeling Kit) or NucGreen Dead as indicated in the results. The nuclear stain was removed by two or three washes in PBS, each lasting 5 min. Stained cells were either imaged directly in PBS or processed for mounting (Fluoroshield +/- DAPI or Prolong Gold Antifade). Images were captured either on an EVOS FL microscope using the 10X and 20X objectives, or on a Leica TCS SP8 confocal laser microscope using an HC PL APO 20X/0.75 dry objective (no. 15506517) or an HC PL APO 20X/0.75 oil-immersion objective (no. 15506343). The LAS X Software from Leica was used to acquire 8-bit confocal images at a resolution of 512 x 512 or 1024 x 1024 pixels, with line averaging set to 8 or 16. For each experiment, the same laser settings were used to image all samples, with the settings optimized to avoid saturation and photobleaching. Images were adjusted for brightness and contrast in FIJI ImageJ, applying the same changes to the entire dataset.

All primary antibodies used for IHC and ICC were first tested individually to confirm specificity and lack of interference in subsequent co-staining experiments.

**RNA processing and RT-qPCR**

Cell lysates were processed for RNA extraction either with the DirectZol Kit or the RNeasy Mini Kit as per the manufacturer's instructions. Live or snap-frozen cells in 12- or 6-well plates were lysed in 500-750µl QIAzol or RLT buffer (supplemented with 10 % (v/v) β-mercaptoethanol). The final RNA was quantified on a NanoDrop ND-1000 and quality assessed by agarose gel electrophoresis. Samples showing signs of RNA degradation were excluded from further analysis. The remaining samples were diluted to 20-100 ng/µl and 200-1,000 ng used for reverse transcription (RT) with the High-Capacity cDNA Reverse Transcription Kit according to the manufacturer's instructions.

For downstream quantitative PCR (qPCR), the cDNA samples were diluted to a final concentration of 1-2.5 ng/µl. Duplicate qPCR reactions were set up in 384-well plates using 2-5 ng/µl cDNA per reaction and SYBR Green chemistry. The samples and the appropriate lineage-specific cDNA standards were run on a Quant Studio 7 Flex Real-Time PCR System (Thermo Fisher Scientific) using the default settings for SYBR reagents, with melting curve acquisition and standard run properties. The thermal conditions were as follows: 2 min at 50 °C; 10 min at 95 °C; 40 cycles at 95 °C for 15 s and 60 °C for 1 min; all with a ramping rate of 1.6 °C/s. Following data acquisition, a mean Ct value was calculated for each sample by taking the average of the technical replicates. Samples were re-run when the technical replicates yielded discordant Ct values (> 0.6 Ct difference), except in cases where this could be attributed to low expression of the targeted gene (e.g. differentiation genes in iPSC samples and stemness genes in differentiated samples). The relative expression value of each gene was determined from the corresponding cDNA standard curve (70). Gene expression values were set to 0 if the Ct values were below the limit-of-detection (LOD) of the standard curve, or within 5 Ct values from the corresponding -RT control(s) when using non-intron-spanning primers. The expression values of individual genes were normalized to the corresponding *TBP* (TATA-binding protein) values, following confirmation that *TBP* expression did not change across conditions. Within each experimental replicate, the final gene expression values were scaled internally to the mean of an appropriate condition to make results comparable across individual experiments.

**TaqMan hPSC Scorecards**

TaqMan hPSC Scorecards (384-well) were used according to the manufacturer's protocol and run on the Quant Studio 7 Flex system. The Ct values were linearized by applying antilog2, based on the assumption of 100 % amplification efficiency. Genes were excluded from analysis if their expression was low (Ct ≥ 30) across all samples. The seven control gene assays were used to calculate a control geometric mean, which was used for normalization of the remaining genes.

**Western blotting**

For protein extraction, adherent cells were lysed in 100-150 µl (per 6-well) ice-cold protein lysis buffer containing: 50 mM HEPES, 150 mM NaCl, 1.5 mM MgCl2, 10 % (v/v) glycerol, 1 % (v/v) TritonX-100, 1 mM EGTA, 100 mM NaF, 10 mM $Na_4P_2O_7$, 2 mM $Na_3VO_4$ (added fresh), 1X EDTA-free protease inhibitor tablet (added fresh), 1X PhosStop tablet (added fresh). Protein concentrations were measured using BioRad's DC protein assay, and all concentrations were adjusted to 1 mg/ml with lysis buffer and LDS sample buffer supplemented with 1X Reducing Agent or 2.5 % (v/v) β-mercaptoethanol. Following denaturation at 70 °C for 10 min, 10 µg of each sample were resolved on NuPAGE 4-12 % gradient gels, using MOPS running buffer supplemented with NuPAGE antioxidant. Next, proteins were transferred to nitrocellulose membranes on an iBlot device (Thermo Fisher Scientific) using program 3 (P3; 20 V) for 7-10 min depending on protein size. Following transfer, the gels were stained with Coomassie (EZ Blue) to assess transfer efficiency and loading accuracy. The nitrocellulose membranes were blocked for 60-90 min in 3 % (v/v) BSA in TBS/T. This was followed by overnight incubation (4 °C) in primary antibody diluted in blocking buffer, up to eight washes in TBS/T and room temperature incubation in horseradish peroxidase (HRP)-conjugated secondary antibodies (diluted 1:10,000) for 1-2 h. The final blots were detected by enhanced chemiluminescence (ECL), using the Immobilon Western Chemiluminescent HRP Substrate and a chemiluminescence imaging analyzer (LAS4000 mini, Fujifilm). Images were processed for cropping and contrast adjustments using FIJI ImageJ (https://imagej.nih.gov/ij/). Densitometry analyses of selected blots were carried out in ImageStudioLite (LI-COR) Version 5.2.5, using the "median" method for background correction.

**Annexin V apoptosis assay**

Apoptosis was assessed with an Annexin V-Cy5 Apoptosis Staining Kit according to a modified version of the manufacturer's protocol. To limit apoptosis due to enzymatic dissociation, the cells were detached with ReLeSR for 15 min prior to collection. The conditioned growth medium and all PBS washes were collected along with the dissociated cells. All samples were centrifuged at 4 °C and 200g for 3 min and the supernatant discarded. Each cell pellet was resuspended in 3 ml ice-cold PBS, followed by repeated centrifugation. The final pellet was resuspended in 250 µl ice-cold 1X Binding Buffer and transferred directly to a FACS tube with a cell strainer lid, allowing the cell suspension to pass through by capillary action. Next, 2.5 µl Annexin Stain were added to each sample, followed by a 20-min incubation on ice prior to fluorescence acquisition. A spare cell suspension was left unstained and used as negative control. A suspension of dead cells was used as positive control. Fluorescence measurements were acquired on a BD Accuri C6 Plus flow cytometer (BD Biosciences) at 675/25 nm with medium flow (35 µl/min) and a core size of 16 µm. At least 50,000 events were acquired per sample. The BD Accuri software was used for gating, ensuring that cell doublets were excluded and that dead, live apoptotic and healthy live cells could be separated based on cell size and Annexin V staining.

#### Study design and replication

A power analysis was carried out in advance of the RNA sequencing analysis, using the method described in (79). According to this analysis, three biological replicates would yield a power of 50 % to detect differentially expressed genes between two genotypes, with a minimum fold-change difference of 1.8 and assuming that 70 % of all genes remain unaltered. This was deemed sufficient for the purpose of this experiment. The RNAseq dataset was subjected to differential expression analysis as described in (72), with FDR $\leq$ 0.05.

"Biological replicates", denoted *N*, refer to independent iPSC clones. Technical replicates refer to multiple cultures of the same iPSC clone, and experimental replicates refer to repeats of the same experiment on different occasions. Additional experimental replicates were included with assays characterized by high internal variability such as spontaneous EB differentiation. The exact number of each type of replicates is stated in the relevant figure legends. Randomization was not required unless in experiments with multiple treatment regiments (e.g. GF stimulations) across different genotypes; in these cases, experimental bias was minimized by treatment and genotype randomization across independent experimental replicates.
