## Supplementary material for "Oncogenic *PIK3CA* promotes cellular stemness in an allele dose-dependent manner"

### Key Resources

| Cell line | Source | Identifier |
|---|---|---|
| WTC11 male parental iPSC line | Coriell Institute | GM25256 |
| WTC11-derived male wild-type post-CRISPR editing | This paper | N/A |
| WTC11-derived male *PIK3CA*$^{WT/H1047R}$ | This paper (CRISPR knock-in) | N/A |
| WTC11-derived male *PIK3CA*$^{H1047R/H1047R}$ | This paper (CRISPR knock-in) | N/A |
| M98-WT female iPSC | This paper (reprogramming of patient-derived fibroblasts) | N/A |
| M98-E418K (*PIK3CA*$^{WT/E418K}$) female iPSC | This paper (reprogramming of patient-derived fibroblasts) | N/A |

**Cell lines**

| Item | Catalogue no. | Vendor | Application |
|---|---|---|---|
| 10 % formaldehyde, Ultrapure, EM- grade, methanol free, | 4018 | PolySciences | Imaging |
| 100 ml DMSO, sterile-filtered | D2650 | Sigma | Cells |
| 2.5 % R.T.U. Horse Serum Blocking (IHC) | S2012 | Vector Laboratories | Imaging |
| 2X SYBR Green PCR Master Mix | 4309155 | Thermo Fisher Scientific | Molbio |
| 2X TaqMan Universal PCR Master Mix | 4318157 | Thermo Fisher Scientific | Molbio |
| Activin A for endoderm/mesoderm EB studies | In-house | Cambridge LRM | Cells |
| Advanced RPMI 1640 | 12633-012 | Thermo Fisher Scientific | Cells |
| Agarose (ultapure) | 16500 | Sigma | Molbio |
| Agencourt CleanSEQ-Dye Termina- tor Removal | A29151 | Beckman Coulter | Molbio |
| AggreWell 400, 24-well | 34411 | Stem Cell Technologies | Cells |
| AggreWell rinsing solution | 7010 | Stem Cell Technologies | Cells |
| Annexin V-Cy5 Apoptosis Staining/Detection Kit | ab14150 | Abcam | Imaging |
| Basic FGF2 for endoderm/mesoderm EB studies | In-house | Cambridge LRM | Cells |
| BigDye Terminator v3.1 Cycle Sequencing Kit | 4337455 | Thermo Fisher Scientific | Molbio |
| Bioanalyzer DNA 12000 Kit | 5067-1508 | Agilent | MolBio |
| Bioanalyzer RNA 6000 Nano Kit | 5067-1511 | Agilent | Molbio |
| BmgBI RE | R0628L | NEB | Molbio |
| BMP4 for endoderm/mesoderm EB studies | In-house | Cambridge LRM | Cells |
| BSA-100g (for blocking) | A7906-100g | Sigma | Protein |

| | | | |
|---|---|---|---|
| BYL719 | B9700-1mg | Cambridge Bioscience | Cells |
| CHIR99021 | 04-0004 | Stemgent | Cells |
| cOmplete ULTRA Tablets, Mini, EasyPack, PhosStop | 5892970001 | Sigma | Protein |
| cOmplete, EDTA-free Protease Inhibitor Cocktail (tablets) | 5056489001 | Sigma | Protein |
| CytoPainter Phalloidin-iFluor 555 Reagent | ab176756 | Abcam | Imaging |
| DAKO antigen retrieval solution pH = 6 (10X) | S2369 | DAKO | Imaging |
| DAKO antigen retrieval solution pH = 9 (10X) | S2367 | DAKO | Imaging |
| DAKO wash buffer 10X | S3006 | DAKO | Imaging |
| DC Protein Assay Reagent A | 5000113 | Biorad | Protein |
| DC Protein Assay Reagent B | 5000114 | Biorad | Protein |
| DC Protein Assay Reagent S | 5000115 | Biorad | Protein |
| Direct-zol RNA Miniprep Kit | R2051 | ZymoResearch | Molbio |
| DMEM/F12 with L-Glutamine, HEPES 500 ml | 11330-032 | Thermo Fisher Scientific | Cells |
| DMEM/F12 without L-Glutamine, HEPES 500 ml | D6421 | Sigma | Cells |
| EndoFree Plasmid Maxi Kit (10) | 12362 | Qiagen | Molbio |
| Essential 6 Medium | A1516401 | Thermo Fisher Scientific | Cells |
| Essential 8 Flex Medium Kit | A2858501 | Thermo Fisher Scientific | Cells |
| Everolimus | HY-10218 | MedChem Express | Cells |
| Exonuclease I | M0293L | NEB | Molbio |
| EZ Blue (Coomassie reagent) | G1041 | Sigma | Protein |
| FACS tubes with cell strainer | 352235 | Corning | Cells/imaging |
| Fetal Bovine Serum (FBS) | SV30180.03 | HyClone | Cells |
| Fluoroshield with DAPI | F6057 | Sigma | Imaging |
| Fluoroshield without DAPI | F6192 | Sigma | Imaging |

| | | | |
|---|---|---|---|
| Geltrex LDEV Free hESC Quality 5 ml | A1413302 | Thermo Fisher Scientific | Cells |
| GeneScan-600 LIZ Size Standard | 4366589 | Applied Biosystems | Molbio |
| GlutaMAX | 35050061 | Thermo Fisher Scientific | Cells |
| Glycerol | 15514-011 | Sigma | Various |
| GoTaq MasterMix (2X) | M7122 | Promega | Molbio |
| HEPES Buffer | H0887 | Sigma | Protein |
| HiDi Formamide 25 ml | 4211320 | Thermo Fisher Scientific | Molbio |
| High-Capacity cDNA Reverse Transcription Kit | 4368814 | Thermo Fisher Scientific | Molbio |
| HiSpeed Plasmid Maxi Kit | 12362 | Qiagen | Molbio |
| Hoechst (Bisbenzimide H 33342 for fluorescence) | 14533-100mg | Sigma | Imaging |
| Horse Serum Blocking (ICC) | 30-0702 | PAN-Biotech | Imaging |
| iBlot transfer stacks regular | IB301001 | Thermo Fisher Scientific | Protein |
| Immobilon Western Chemiluminescent HRP Substrate | WBKLS0500 | Merck Millipore | Protein |
| Ingenio Cuvettes 50 pk | MIR50121 | Cambridge Bioscience | Nucleofection |
| KaryoMAX Colcemid Solution in PBS | 15212012 | Thermo Fisher Scientific | Cells |
| Matrigel, hESC-qualified | 354277 | Corning | Cells |
| mFreSR freezing reagent 50 ml | 5855 | Stem Cell Technologies | Cells |
| MinElute PCR Purification Kit 50 rxn | 28004 | Qiagen | Molbio |
| Nuclear Select FX Labelong kit | S33025 | Thermo Fisher Scientific | Imaging |
| Nunclon Sphera 60 mm dish (21.5 cm2 ) | 174944 | Thermo Fisher Scientific | Cells |

| | | | |
|---|---|---|---|
| NuPAGE 4-12% BT midi 12+2 well PAGE gels | WG14001BOX | Thermo Fisher Scientific | Protein |
| NuPAGE Antioxidant | NP0005 | Thermo Fisher Scientific | Protein |
| NuPAGE LDS Sample Buffer (4X) | NP0007 | Thermo Fisher Scientific | Protein |
| NuPAGE MOPS SDS Running Buffer 20X | NP0001 | Thermo Fisher Scientific | Protein |
| NuPAGE Sample Reducing Agent (10X) | NP0009 | Thermo Fisher Scientific | Protein |
| OneShot Stbl3 Chemically Competent Cells | C7373-03 | Thermo Fisher Scientific | Molbio |
| PBS | D8537 | Sigma | Cells |
| Phusion High-Fidelity DNA Polymerase | M0530L | NEB | Molbio |
| Precision Plus Dual Color Protein Standard | 161-0374 | Biorad | Protein |
| Prolong Gold Antifade reagent | P36930 | Thermo Fisher Scientific | Imaging |
| PSC Cryopreservation Kit | A2644601 | Thermo Fisher Scientific | Cells |
| Q5 High-Fidelity DNA Polymerase | M0491L | NEB | Molbio |
| QIAamp DNA Micro Kit (50) | 56304 | Qiagen | Molbio |
| QIAzol Lysis Reagent | 79306 | Qiagen | Molbio |
| QuickExtract DNA Extraction Solution | QE09050 | Cambridge Bioscience | Molbio |
| Ready Probes Cell Viability Imaging Kit, Blue/Green | R37609 | Thermo Fisher Scientific | Imaging |
| Recombinant human EGF 100 µg | AF-100-15 | PeproTech | Cells |
| Recombinant human FGF2-basic (for GF stimulations) | 100-18B | PeproTech | Cells |
| Recombinant human IGFI 100 µg | 100-11 | PeproTech | Cells |
| ReLeSR 100 ml | 5872 | Stem Cell Technologies | Cells |

| | | | |
|---|---|---|---|
| RevitaCell Suppplement 5 ml | A2644501 | Thermo Fisher Scientific | Cells |
| RLT Buffer | 79216 | Qiagen | Molbio |
| RNeasy Mini Kit | 74104 | Qiagen | Molbio |
| SAP (Shrimp Alkaline Phosphatase) | GEE70092Z | Sigma | Molbio |
| Secondary antibody (ICC), Donkey anti-Goat IgG (H+L) Cross-Adsorbed Secondary Antibody, Alexa Fluor 647 | A21447 | Thermo Fisher Scientific | Imaging |
| Secondary antibody (ICC+IHC), Goat anti-Mouse IgG (H+L) Highly Cross-Adsorbed Secondary Antibody, Alexa Fluor 488 | A11029 | Thermo Fisher Scientific | Imaging |
| Secondary antibody (ICC), Goat anti-Rabbit IgG (H+L) Highly Cross-Adsorbed Secondary Antibody, Alexa Fluor 594 | A11037 | Thermo Fisher Scientific | Imaging |
| Secondary antibody (WB), Goat anti-Rabbit IgG HRP-linked Antibody | 7074S | CST | Protein |
| Secondary antibody (WB), Horse anti-Mouse IgG HRP-linked Antibody | 7076S | CST | Protein |
| StemPro Accutase | A110501 | Thermo Fisher Scientific | Cells |
| SuperScript IV Fist Strand Synthesis System | 18091050 | Thermo Fisher Scientific | Molbio |
| TaqMan® hPSC ScorecardTM Panel, 384-well | A15870 | Thermo Fisher Scientific | Molbio |
| TeSR-E8 Medium Kit | 5940 | Stem Cell Technologies | Cells |
| TGX221 | S1169 | SelleckChem | Cells |
| TrueSeq mRNA library kit | 20020594 | Illumina | Molbio |

| | | | |
|---|---|---|---|
| TruSeq RNA Single Indexes Set A | 20020492 | Illumina | Molbio |
| TrypLe Express Enzyme (1X), phe- nol red | 12605010 | Thermo Fisher Scientific | Cells |
| Wizard Plus SV Miniprep Kit | A1330 | Promega | Molbio |
| β-mercaptoethanol | M7522 | Sigma | Protein |
| μ-Dish 35 mm, low | 80136 | Ibidi | Imaging |
| μ-Slide 4 Well | 80426 | Ibidi | Imaging |

**Materials list.** ICC, immunocytochemistry; IHC, immunohistochemistry; Molbio, molecular biology; WB, Western blotting.

| Component | Sequence |
| --- | --- |
| sgRNA-Fb | ATGAATGATGCACATCATGG |
| HDR001 | TAGCCTTAGATAAAACTGAGCAAGAGGCTTTGGAGTATTTCATGAAACAAA<br>TGAACGACGCACGTCATGGTGGCTGGACAACAAAAATGGATTGGATCTTCC<br>ACACAATTAAACAGCATGCATTGAACTGAAAAGATAACTGAGAAAATG |
| HDR002 | TAGCCTTAGATAAAACTGAGCAAGAGGCTTTGGAGTATTTCATGAAACAAATG<br>AACGACGCACATCATGGTGGCTGGACAACAAAAATGGATTGGATCTTCCACAC<br>AATTAAACAGCATGCATTGAACTGAAAAGATAACTGAGAAAATG |

**Nucleic acid sequences of sgRNA-Fb and the two HDR templates used to target *PIK3CA* exon 21.** All sequences are provided in the 5’-to-3’ direction.

| **Primer ID** | **Sequence** | **Description** |
|---|---|---|
| PR005 | CAGCATGCCAATCTCTTCAT | Sequencing of *PIK3CA* exon 21 (H1047R) |
| PR006 | ATGCTGTTCATGGATTGTGC | Sequencing of *PIK3CA* exon 21 (H1047R) |
| PR009-F | FAM-TGATGCTTGGCTCTGGAATG | RFLP for *PIK3CA* exon 21; annealing T = 56 °C for 20 or 30 s. Use with PR010-R. |
| PR010-R | GGTCTTTGCCTGCTGAGAGT | RFLP for *PIK3CA* exon 21; annealing T = 56 °C for 20 or 30 s. Use with PR009-F. |
| Off-target-1F | TATGGCTCCTCTGATTGCACC | Off-target sequencing |
| Off-target-1R | CAGGCACCCATTTCTGGTAA | Off-target sequencing |
| Off-target-2F | GAGCCACGGACTTCCAGATAG | Off-target sequencing |
| Off-target-2R | CTAGCACAGGGCCACACAAAT | Off-target sequencing |
| Off-target-3F | GTGAAGTGAGGCGTGATGGG | Off-target sequencing |
| Off-target-3R | AGCAGAAGACCTCAACCATCC | Off-target sequencing |
| Off-target-4F-nest | CTACCTCCAAGCACATGGCA | Off-target sequencing |

| | | |
|---|---|---|
| Off-target-4R-nest-inner | TGTGCCTCAATGTGCAAAGC | Off-target sequencing |
| Off-target-4R-nest-outer | TGAGCCCTTACCCTGTAGGG | Off-target sequencing |
| Off-target-5F-1 | CAGAAATTTAGGGAAGGTGAATGG | Off-target sequencing |
| Off-target-5F-2 | AGTTATCTGTTGTACAACATAGTGCC | Off-target sequencing |
| Off-target-5R-1 | GATGTGTAAGGGAGAGCTGGTA | Off-target sequencing |
| Off-target-5R-2 | CAGGATCAGATGGATTCACAACA | Off-target sequencing |
| Off-target-6F | GCCACACTCCACTGGTAGAC | Off-target sequencing |
| Off-target-6R | CAGTGTGCGTCTTCCCTTCT | Off-target sequencing |
| Off-target-7F | TTCTGGACCCCACCTCAGTA | Off-target sequencing |
| Off-target-7R | CCTTGAACATAGCCCCACCA | Off-target sequencing |
| Off-target-8F | TGGCTGGTAAGCACTCTCCT | Off-target sequencing |
| Off-target-8R | CCCCTGGCCTTTACGCTAAC | Off-target sequencing |
| Off-target-9F | TGCAGACTGAGAGTGCATGT | Off-target sequencing |
| Off-target-9R | GGCAGCTAGGGTTCTCAGTG | Off-target sequencing |

| | | |
|---|---|---|
| Off-target-10F | ATTGTGGGACCACAGGAACA | Off-target sequencing |
| Off-target-10R | GGGCAAGGAAGCACTCTTGT | Off-target sequencing |
| Off-target-12F | CACATGACAGCTCATGGTGATG | Off-target sequencing |
| Off-target-12R | TGTGGAGGGCAACATACATCT | Off-target sequencing |
| Off-target-13F | CTGATGCTGCCCCTATACCA | Off-target sequencing |
| Off-target-13R | AAACTACCTGACACACTATAAGCA | Off-target sequencing |
| Off-target-14F | CCTTACACTGGCTGACCCTG | Off-target sequencing |
| Off-target-14R | GGGGCCACATCTTACTGCTT | Off-target sequencing |
| Off-target-15F | CTCAAAAGTCGCAGTGGCTC | Off-target sequencing |
| Off-target-15R | GTGACCAAATGGAAGCTGGC | Off-target sequencing |
| Off-target-16F | CTGGAAGAAGGGACGTCAGAG | Off-target sequencing |
| Off-target-16R-2 | AGCAAATGGTACTGTCAAGTCC | Off-target sequencing |
| Off-target-17F | TATGCCTTCAAGACCCGCAA | Off-target sequencing |

| | | |
|---|---|---|
| Off-target-17R | AAGCTTCCGCTTCTAGGCAAG | Off-target sequencing |
| Off-target-18F | CAACCCTCCCTGAGTACGTG | Off-target sequencing |
| Off-target-18R | TGTGGGTCATCGAGGGTCTG | Off-target sequencing |
| PIK3CAex21region-1F | TGCTGTGAAGGAAAATGGAAAGG | Tiled primer matrix, *PIK3CA* exon 21 |
| PIK3CAex21region-2F | TACAACACAGTCCTGACTCTAGC | Tiled primer matrix, *PIK3CA* exon 21 |
| PIK3CAex21region-3F | ACTGAGCAAGAGGCTTTGGA | Tiled primer matrix, *PIK3CA* exon 21 |
| PIK3CAex21region-4F | CGCAAACAGGGTTTGATAGCAC | Tiled primer matrix, *PIK3CA* exon 21 |
| PIK3CAex21region-5F | TGGGGTGGAAGGGACTCTTG | Tiled primer matrix, *PIK3CA* exon 21 |
| PIK3CAex21region-6F | TTGCTCCAAACTGACCAAACT | Tiled primer matrix, *PIK3CA* exon 21 |
| PIK3CAex21region-1R | AGTTTGGTCAGTTTGGAGCAA | Tiled primer matrix, *PIK3CA* exon 21 |
| PIK3CAex21region-2R | CGGTCTTTGCCTGCTGAGA | Tiled primer matrix, *PIK3CA* exon 21 |
| PIK3CAex21region-3R | CAGCCTTTGTTGTGTCCACATT | Tiled primer matrix, *PIK3CA* exon 21 |
| PIK3CAex21region-4R | GACTGCTTCCAAAACTGCAGA | Tiled primer matrix, *PIK3CA* exon 21 |
| | GTATTCAGTTCAATTGCAGAAGGAG | Tiled primer matrix, *PIK3CA* exon 21 |

| | | |
|---|---|---|
| PIK3CAex21region-5R-1 | | |
| PIK3CAex21region-5R-2 | GCTGACCATGCTGCTATGAAC | Tiled primer matrix, *PIK3CA* exon 21 |
| PIK3CAex21region-6R | GTGCTATCAAACCCTGTTTGCG | Tiled primer matrix, *PIK3CA* exon 21 |
| TP53-exon2+3-F-nest-outer | GCTTGGGTTGTGGTGAAACA | *TP53* sequencing |
| TP53-exon2+3-F-nest-inner | GCAGCCATTCTTTTCCTGCTC | *TP53* sequencing |
| TP53-exon2+3-R-nest | GGTGAAAAGAGCAGTCAGAGGA | *TP53* sequencing |
| TP53-exon4-F-nest | AGAGACCTGTGGGAAGCGAA | *TP53* sequencing |
| TP53-exon4-R-nest-outer | CAGGAAGCCAAAGGGTGAAGA | *TP53* sequencing |
| TP53-exon4-R-nest-inner | GGATACGGCCAGGCATTGAA | *TP53* sequencing |
| TP53-exon5+6-F | GAGGTGTAGACGCCAACTCT | *TP53* sequencing |
| TP53-exon 5+6-R | CCACTGACAACCACCCTTAAC | *TP53* sequencing |
| TP53-exon7-F | CTCATCTTGGGCCTGTGTTATCT | *TP53* sequencing |
| TP53-exon7-R | GTGATGAGAGGTGGATGGGTAG | *TP53* sequencing |
| TP53-exon8+9-F | CTACCCATCCACCTCTCATCAC | *TP53* sequencing |

| | | |
|---|---|---|
| TP53-exon8+9-R | ATGCCCCAATTGCAGGTAAAAC | *TP53* sequencing |
| TP53-exon10-F-nest | AATGCATGTTGCTTTTGTACCGTC | *TP53* sequencing |
| TP53-exon10-R-nest-outer | CCCTGGGTTTGGATGTTCTG | *TP53* sequencing |
| TP53-exon10-R-nest-inner | TGACCATGAAGGCAGGATGA | *TP53* sequencing |
| TP53-exon11-F-nest | AGCATTGGTCAGGGAAAAGG | *TP53* sequencing |
| TP53-exon11-R-nest-outer | AAGCAAGGGTTCAAAGACCCA | *TP53* sequencing |
| TP53-exon11-R-nest-inner | CAAAACCCAAAATGGCAGGGG | *TP53* sequencing |

**Supplementary Table 3. Genotyping primers.** F and R denote forward and reverse direction, respectively; all given in the 5'-to-3' direction. "-nest" in the ID indicates that the primer was used in nested PCR reactions, with "outer" and "inner" specifying that the primer was used in the first and second amplification, respectively. Primers used to sequence PIK3CA across codon H1047 required 56 °C during annealing. The annealing temperature for the remaining primers was 60 °C.

| Target | Catalog # (clone if mAb) | Vendor | Species | Molecular weight (kDa) | Dilution | Application |
|---|---|---|---|---|---|---|
| FGFR1 | 9740 (XP D8E4) | CST | Rabbit mAb | 92, 120, 145 | 1:1,000 | Western |
| IGF1Rβ | 9750 (XP D23H3) | CST | Rabbit mAb | 95 (mature), 200 (proform) | 1:1,000 | Western |
| IRβ (C-19) | sc-711 | Santa Cruz | Rabbit pAb | 90 (mature), 200 (proform) | 1:200 | Western |
| p110α | 4249 (C73F8) | CST | Rabbit mAb | 110 | 1:1,000 | Western |
| p110β | 3011 (C33D4) | CST | Rabbit mAb | 110 | 1:1,000 | Western |
| pAKT T308 | 9275 | CST | Rabbit pAb | 60 | 1:1,000 | Western |
| pAKT S473 | 9271 | CST | Rabbit pAb | 60 | 1:1,000 | Western |
| pAKT S473 | 4060 (9DE XP) | CST | Rabbit mAb | 60 | 1:1,000 | Western |
| AKT | 9272 | CST | Rabbit pAb | 60 | 1:1,000 | Western |
| pS6K1 T389 | 9205 | CST | Rabbit pAb | 70, 85 | 1:1,000 | Western |
| pS6 S235/S236 | 2211 | CST | Rabbit pAb | 32 | 1:1,000 | Western |
| S6 | 2217 (5G10) | CST | Rabbit mAb | 32 | 1:1,000 | Western |

| | | | | | | |
|---|---|---|---|---|---|---|
| pERK1/2 T202/Y204, T185/Y187 | 9106 (E10) | CST | Mouse mAb | 42,44 | 1:1,000 | Western |
| pERK1/2 T202/Y204, T185/Y187 | 4377 (197G2) | CST | Rabbit mAb | 42,44 | 1:1,000 | Western |
| pERK1/2 T202/Y204, T185/Y187 | 4370 (D13.14.4E) | CST | Rabbit mAb | 42,44 | 1:1,000 1:2,000 | Western |
| ERK1/2 | 4695 (137F5) | CST | Rabbit mAb | 42,44 | 1:1,000 | Western |

**Primary antibodies used for Western blotting.** CST, Cell Signaling Technology; kDa, kilodalton; mAb, monoclonal antibody; pAb, polyclonal antibody.

| Target | Catalog # (clone if mAb) | Vendor | Species | Conjugate | Dilution | Application |
|---|---|---|---|---|---|---|
| TUBB3 | 801201 (TUJ1) | Biolegend | Mouse | No | 1:500 (2 µg/ml) | ICC |
| T BRACHYURY | AF2085 | R&D | Goat pAb | No | 1:100 (2 µg/ml) | ICC/IHC |
| α -SMA (ACTA2) | A5228 (1A4) | Sigma | Mouse mAb | No | 1:200 (10 µg/ml) | ICC |
| HAND1 | AF3168 | R&D | Goat pAb | No | 1:100 (2 µg/ml) | ICC |
| SOX17 | AF1924 | R&D | Goat pAb | No | 1:200 (1 µg/ml) | ICC |
| FOXA2 (hHNF3β ) | AF2400 | R& D | Goat pAb | No | 1:50 (4 µg/ml) | ICC |
| AFP | sc-8399 (C3) | Santa Cruz | Mouse mAb | No | 1:100 (2 µg/ml) | ICC |
| AFP | 14-9499-82 (1E8) | Thermo Fisher Scientific | Mouse mAb | No | 1:100 (5 µg/ml) | IHC |
| OCT3/4 | sc-5279 (C-10) | Santa Cruz | Mouse mAb | No | 1:100 (2 µg/ml) | ICC/IHC |
| TRA-1-60 | sc-21705 | Santa Cruz | Mouse mAb | No | 1:100 (2 µg/ml) | ICC |
| TRA-1-60 | 14-8863-82 | Thermo Fisher Scientific | Mouse mAb | No | 1:100 (5 µg/ml) | ICC |
| NANOG | AF1997 | R&D | Goat pAb | No | 1:100 (2 µg/ml) | ICC |
| NANOG | PA1-097 | Thermo Fisher Scientific | Rabbit pAb | No | 1:100 (10 µg/ml) | ICC |
| Cyclophilin A | 51418 (D2Y4M) | CST | Rabbit mAb | No | 1:100 (1.3 µg/ml) | IHC |
| Mouse IgG isotype control | I5381 | Sigma | Mouse | No | 1:1975 (2 µg/ml); 1:395 (10 µg/ml) | ICC |

| | | | | | | |
|---|---|---|---|---|---|---|
| Mouse IgG isotype control | 37355 | Abcam | Mouse | No | 1:2500 (2 μg/ml); 1:1000 (5 μg/ml); 1:500 (10 μg/ml) | IHC |
| Rabbit IgG isotype control | 10500C | Thermo Fisher Scientific | Rabbit | No | 1:1500 (2 μg/ml); 1:300 (10 μg/ml) | ICC |
| Goat IgG isotype control | ab37373 | Abcam | Goat | No | 1:1500 (2 μg/ml) | ICC/IHC |

**Primary antibodies used for immunocytochemistry (ICC) and immunohistochemistry (IHC).** CST, Cell Signaling Technology; mAb, monoclonal antibody; pAb, polyclonal antibody.

| **Target** | **Accession ID (RefSeq)** | **Fwd primer** | **Rev primer** | **Amplicon size (bp)** |
|---|---|---|---|---|
| *POU5F1 (OCT3/4)* | NM_002701.5 | TGTACTCCTCGGTCCCTTTC | TCCAGGTTTTCTTTCCCTAGC | 150 |
| *NANOG* | NM_024865.3 | CAGTCTGGACACTGGCTGAA | CTCGCTGATTAGGCTCCAAC | 149 |
| *FGF4* | NM_002007.2 | CCAACAACTACAACGCCTACGA | CCCTTCTTGGTCTTCCCATTCT | 82 |
| *GDF3* | NM_020634.2 | TTTCTCCCAGACCAAGGTTTC | TCCCTTTCTTTGATGGCAGA | 107 |
| *NODAL* | NM_018055.4 | CAGTACAACGCCTATCGCTGT | TGCATGGTTGGTCGGATGAAA | 75 |
| *KDR (VEGFR2)* | NM_002253.2 | GTGATCGGAAATGACACTGGAG | CATGTTGGTCACTAACAGAAGCA | 124 |
| *T BRACHYURY* | NM_003181.3 | GCAAAAGCTTTCCTTGATGC | ATGAGGATTTGCAGGTGGAC | 144 |
| *HAND1* | NM_004821.2 | TCCGCAGAAGGGTTAAACAG | CAAGGCTGAAAATGAGACGC | 99 |
| *DKK1* | NM_012242.2 | ACCAGCTATCCAAATGCAG | TCACAGGGGAGTTCCATAAA | 181 |
| *SOX17* | NM_022454.3 | CGCACGGAATTTGAACAGTA | CAGTAATATACCGCGGAGCTG | 149 |
| *FOXA2* | NM_021784.4 | GGAGCAGCTACTATGCAGAGC | CGTGTTCATGCCGTTCATCC | 83 |
| *CXCR4* | NM_003467.2 | GCCTTATCCTGCCTGGTATTGTC | GCGAAGAAAGCCAGGATGAGGAT | 130 |

| | | | | |
|---|---|---|---|---|
| *AFP* | NM_001134.2 | GCTTTGCTGAAGAGGGACAA | ACACCGAATGAAAGACTCGT | 105 |
| *HNF4A* | NM_001287182.1 | ATCGCAGATGTGTGTGAGTC | GCAGAAAGCTGGGATGTACT | 78 |
| *PAX6* | NM_001604.5 | AACGATAACATACCAAGCGTGT | GGTCTGCCCGTTCAACATC | 120 |
| *OTX2* | NM_021728.3 | CAAAGTGAGACCTGCCAAAAAGA | TGGACAAGGGATCTGACAGTG | 179 |
| *TBP* | NM_003194.4 | TAATCCCAAGCGGTTTGC | TAGCTGGAAAACCCAACTTCT | 170 |

**SYBR Green qPCR primer.** Bp, base pairs; Fwd, forward; Rev; reverse. Note that the lineage specification is only guiding as most genes are also expressed in other lineages. Only a single accession ID is given due to space constraints, but several of the primers detect additional splice isoforms (*TBP*, *POU5F1*, *NANOG*, *NODAL*, *FOXA2*, *CXCR4*, *HNF4A*, *PAX6, OTX2*).

| Data | Source | Identifier |
|---|---|---|
| Raw and analysed data | This paper | Open Science Framework, doi: xxxxx (to be provided upon acceptance) |
| Protein-coding RNA sequencing | This paper | GEOXXXXX (to be provided upon acceptance) |

**Deposited data**
